# Non-parametric Bayesian density estimation for biological sequence space with applications to pre-mRNA splicing and the karyotypic diversity of human cancer

**DOI:** 10.1101/2020.11.25.399253

**Authors:** Wei-Chia Chen, Juannan Zhou, Jason M Sheltzer, Justin B Kinney, David M McCandlish

## Abstract

Density estimation in sequence space is a fundamental problem in machine learning that is of great importance in computational biology. Due to the discrete nature and large dimensionality of sequence space, how best to estimate such probability distributions from a sample of observed sequences remains unclear. One common strategy for addressing this problem is to estimate the probability distribution using maximum entropy, i.e. calculating point estimates for some set of correlations based on the observed sequences and predicting the probability distribution that is as uniform as possible while still matching these point estimates. Building on recent advances in Bayesian field-theoretic density estimation, we present a generalization of this maximum entropy approach that provides greater expressivity in regions of sequence space where data is plentiful while still maintaining a conservative maximum entropy character in regions of sequence space where data is sparse or absent. In particular, we define a family of priors for probability distributions over sequence space with a single hyper-parameter that controls the expected magnitude of higher-order correlations. This family of priors then results in a corresponding one-dimensional family of maximum a posteriori estimates that interpolate smoothly between the maximum entropy estimate and the observed sample frequencies. To demonstrate the power of this method, we use it to explore the high-dimensional geometry of the distribution of 5′ splice sites found in the human genome and to understand the accumulation of chromosomal abnormalities during cancer progression.

## Introduction

Biological data is often discrete and combinatorial. We observe, for instance, some collection of macromolecular sequences that take the form of a string of nucleotides or amino acids. Or we make a multi-channel neural recording, resulting in a collection of strings composed of 0s and 1s corresponding to the set of neurons that are firing at each instant in time. A natural question given a collection of such strings is whether we can estimate the probability distribution that these sequences are drawn from [1, 2, 3, 4, 5].

Estimating such a probability distribution can be challenging because the number of possible sequences grows exponentially in sequence length, and even if the subset of biologically active or relevant sequences is small compared to the entirety of the space, this biologically relevant subset can still easily contain thousands of sequences. As a result, estimating the frequency of each possible sequence becomes impractical and we require some prior or set of simplifying assumptions in order to make progress.

Among the most common simplifying assumptions is that the true distribution takes the form of a maximum entropy distribution, defined as the most uniform (i.e. highest entropy) distribution compatible with certain summary statistics of the sample [6]. In a typical case, these summary statistics are taken to be the frequency of each possible letter at each position, which produces the ubiquitous matrix model or position weight matrix that represents a probability distribution of sequences in terms of independent probability distributions at each position in the sequence [7, 8]. However, it is also common to attempt to capture correlations between positions, which results in pairwise maximum entropy models also known as Potts models [1, 9, 10, 11, 12, 13, 14, 15, 16, 17, 18]. Such pairwise interaction models have seen great success in a variety of applications, including identifying functional elements [9], predicting residues or positions that contact each other or interact [14, 19, 20], and predicting the effects of mutations [21].

Here we provide a generalization of these maximum entropy models that can achieve greater expressivity in well-sampled, high probability regions of sequence space while still providing parsimonious density estimates in low probability regions of sequence space, where data is by necessity sparse or absent. We do this by deriving a one-parameter family of Bayesian priors for probability distributions over sequence space capable of capturing correlations of all orders, with the single hyper-parameter controlling the expected extent of the departure from the corresponding maximum entropy model. The resulting family of maximum a posteriori estimates all match the same moments as the corresponding maximum entropy model, and includes the maximum entropy model and the histogram of observed frequencies as limiting cases. Importantly, in non-limiting cases these models produce estimates whose character is similar to the histogram of observed frequencies where the data is able to overwhelm the prior, but which exhibit the smooth behavior typical of a maximum entropy model in poorly sampled regions of sequence space. At a more technical level, the method we propose is the discrete multivariate analog of recently developed field-theoretic techniques [22, 23, 24, 25, 26, 27, 28] for real-valued random variables known as DEFT (Density Estimation using Field Theory, [28]), and we therefore refer to our method as SeqDEFT.

In what follows, we describe the formal features of our method and then apply the method to explore two datasets. The first of these datasets is the collection of all annotated 5’ pre-mRNA splice sites in the human genome (RNA sequences of length 9). Because of the relatively large number of annotated splice sites (305,106) compared to the number of possible sequences (4^9^ = 262,144) we use this dataset to explore the behavior of our method on complex distributions as increasing amounts of data are available. For the second dataset, we consider the distribution of chromosomal copy number abnormalities observed across human cancers [29]. Here, there are far fewer observations (10,522 samples) than there are possible karyotypic patterns (2^22^ = 4,194,304 under the simplest scoring where each of the 22 autosomes is scored as being either euploid or aneuploid). However, our method is still able to recover the signatures of several complex aneuploid states corresponding to sets of multiple chromosomes that are frequently altered together. We also provide visualizations and a qualitative exploration of the major features of the inferred high-dimensional probability distributions for each of these two datasets, with an emphasis on understanding the biological basis for the complex, multi-modal structure of these distributions.

## Results

We consider probability distributions defined on the Hamming graph of sequences with length *£* and *α* alleles per position, where two sequences are adjacent if they differ in exactly one position. To define a probability distribution *Q* on this graph, we first define a field *ϕ* on the graph and then let the probability of drawing sequence *i* be given by

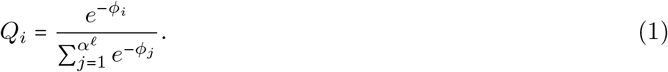

Thus, we can define a prior over probability distributions by imposing a prior on this field *ϕ*.

### Extending the independent model

The maximum entropy model based on the marginal frequencies of the alleles at each position is equivalent to making an independent draw of the allele for each position from the observed marginal frequencies. For concreteness, we will derive our method for this basic form of maximum entropy model before turning to the general case.

Our overall strategy is to think geometrically about *ϕ* as a function on the Hamming graph. In particular, let us consider the behavior of *ϕ* on one particular “face” of the Hamming graph, which we can think of in terms of wild-type sequence *i*, two single mutants *j* and *k*, and a double mutant *m*. Moreover, if we draw a sequence from this face, then we can capture the conditional association between these two mutations in terms of the log odds ratio:

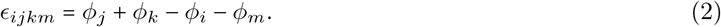

For the special case of the independent model the values of *ϕ* are additive such that double mutant *ϕ*_*m*_ is given by *ϕ* evaluated at the wild-type, plus the effects of each of the two single mutants on *ϕ*, i.e.

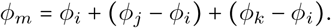

Rearranging this expression, it is easy to see that this implies that for the independent sites maximum entropy model the conditional log odds ratio *E*_*ijkm*_ is zero for every face in the Hamming graph.

To build a model that allows deviations from the maximum entropy assumption but tends to make these deviations be small, we can thus construct a prior on functions *ϕ* where the probability of *ϕ* is a function of the average squared conditional log odds ratio 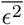, which quantifies the average local deviation from independence where this average is taken over all the faces of the Hamming graph. In fact, it is possible to derive a simple formula for 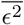 in terms of the graph Laplacian *L* of the Hamming graph, where *L* is defined as:

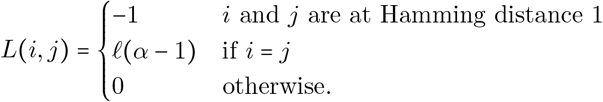

In particular, if we let Δ = (*L*^2^ − *αL*)/2 and let 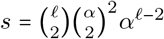 be the number of faces, then this average is given by the quadratic form 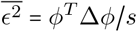 [30].

Using this expression for the mean squared log odds ratio, we can then define a family of Gaussian priors on *ϕ* with probability density

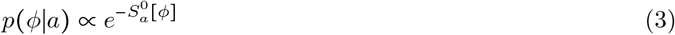

in terms of the action

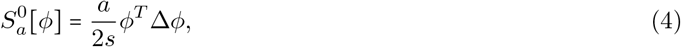

where *a* is a hyper-parameter that controls the expected magnitude of deviations from independence, with larger values of *a* producing smaller deviations. In particular, in the SI Appendix we show that the expectation of 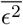 under the prior is given by rank (Δ)/*a*, where in this case rank (Δ) = *α*^*l*^ – (*α* – 1)*l* – 1 (see SI Appendix). We also show that this prior is equivalent to independently drawing the value of *ϵ* for each face in the Hamming graph from a zero-mean Gaussian with variance *s/a* and then conditioning on these local correlations being globally consistent with each other (c.f., [31]) in the sense of being simultaneously realizable by some choice of *ϕ*.

Given this prior distribution and a sample of size *N* with realized frequencies given by the vector *R*, we find that the posterior distribution has 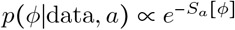 where the posterior action *S*_*a*_ [*ϕ*] is given by

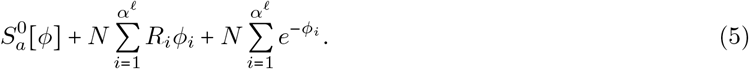

The maximum a posteriori (MAP) estimate is then found by minimizing this action. We show in the SI appendix that the MAP estimate approaches the maximum entropy solution in the limit *a* → ∞, approaches the empirically observed frequencies in the limit as *a* → 0, and matches the same moments as the maximum entropy solution for all values of *a*, where this behavior mirrors existing results on field-theoretic Bayesian density estimation [27] for continuous one-dimensional random variables but instead applies to estimating probability distributions in sequence space.

### Extending pairwise and higher-order maximum entropy models

So far for concreteness we have concentrated on providing a non-parametric extension to the independent model, which is the maximum entropy model that matches the observed frequencies of the alleles at each position. However, we can generalize the above approach to provide analogous results for pairwise and higher-order [32] maximum entropy models by making a corresponding change to the prior.

In particular, let us consider the maximum entropy model that matches the moments of our observations up to order *P* − 1. For *P* = 2 we have the independent sites model that matches the site-specific allele frequencies, and each possible mutation (parallel edges of the Hamming graph) has a constant effect on *ϕ* so that the conditional log odds ratios defined on each face of the Hamming graph is zero. In the proceeding section, we constructed a relaxation of this maximum entropy model by considering the average squared conditional log odds ratio, where this average is taken over all faces in the Hamming graph.

To extend this same idea to *P* = 3, which is the pairwise interaction model that matches the site-specific allele frequencies and pairwise correlations between sites, we note that this model has constant conditional log odds ratios for faces defined by the same pair of mutations (i.e. parallel faces) and hence the difference between conditional log odds ratios on adjacent parallel faces of the Hamming graph are all 0. These adjacent parallel faces form 3-faces or cubes, and hence we can extend this model using a prior that produces an expected squared difference between the conditional log odds ratios on parallel faces of these cubes. More generally for a maximum entropy model that matches the first *P* − 1 moments, our extension is based on defining a prior in terms of the expected mean squared deviation from the local behavior implied by the corresponding maximum entropy assumption, where the average is taken over all sub *P* -faces of the Hamming graph. More formally, to define this prior, consider the operator:

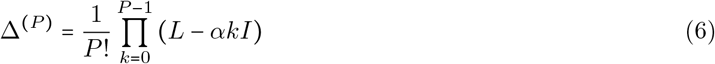

where *I* is the identity matrix. Then define the prior action to be

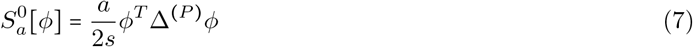

where 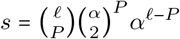 is the number of *P* -dimensional faces of the Hamming graph. Under this prior, the MAP estimate always matches the first *P* − 1 moments of the observations, the limit as *a* → ∞ results in the maximum entropy model and the limit as *a* → 0 results in the empirically observed distribution, rank (Δ(^(*P*)^)/*a* gives the expected mean squared conditional *P* -association under the prior, and we can likewise construct this prior by drawing the conditional *P* -association for each *P* -face from a zero-mean normal distribution with variance *s*/*a* (see SI Appendix for details).

### Practical implementation

There are several issues that need to be addressed to produce a practical implementation of SeqDEFT. First, due to the form of Equation 6, most of the key aspects of the necessary computations can be rewritten in terms of repeated matrix multiplications by the extremely large matrix *L*. Fortunately, *L* is also very sparse, and we can exploit this sparsity to conduct our calculations for sequence spaces containing up to several million genotypes. Another issue is how to choose the value of the hyper-parameter *a*. Here we choose this value by maximizing the *k*-fold cross-validated log-likelihood with *k* = 5, and refer to the resulting optimal value as *a*^∗^. However, such cross-validation is computationally demanding as one has to repeat the same computations *k* + 1 times. We overcome this difficulty by parallelizing the computations on a cluster. In the SI Appendix we describe an alternative approach for finding *a*^∗^ (for small sequence spaces) using the evidence ratio. Finally, sampling from the posterior distribution in such high-dimensional sequence spaces is very challenging. A relatively efficient method of posterior sampling in such circumstances is the Hamiltonian Monte Carlo [33] which employs Hamiltonian dynamics to help the imaginary particle probe the probability landscape; see the SI Appendix for details on our implementation.

### Distribution of human 5’ splice sites

In most eukaryotes, the sequence of the final processed form of an mRNA transcript is not encoded contiguously in the genome but rather appears as several discrete segments known as exons that are separated by other DNA segments known as introns. During transcription, both the intronic and exonic DNA are transcribed into RNA, after which the intronic RNA is removed in a process known as pre-mRNA splicing [34]. To demonstrate the characteristics of our SeqDEFT method, we first considered 5’ splice sites, the RNA sequences at the boundary between each intron and its upstream exon [35]. Because there are hundreds of thousands of such sites in the human genome, 5’ splice sites provide a relatively well-sampled model system for understanding the complexity and geometry of high-dimensional distributions in sequence space, as well as an opportunity to sub-sample real data in order to investigate the performance of our method when less data is available. In addition, modeling and identification of splice sites was one of the early successes of maximum entropy models with pairwise interactions and remains a common method for scoring splice sites [9].

For each annotated intron in the human genome, we extracted the last 3 positions of the upstream exon (which are typically labeled as −3,-2,-1) and the first 6 positions of intron itself (labeled +1 through +6). In total this resulted in a collection of 305,106 9-nucleotide sequences, which we modeled using SeqDEFT. To understand the qualitative behavior of SeqDEFT on datasets of different sizes, we further performed a rarefaction analysis, where we trained models on 25%, 5% or 1% of the data. In the first row of Figure 1, we see that at very low sampling, the SeqDEFT MAP estimate *Q*^∗^ behaves very similarly to the independent positions maximum entropy model, but becomes substantially different as the amount of data increases. The second row of Figure 1 compares the SeqDEFT estimate from the subsampled data to the estimate using the full dataset and we see that the SeqDEFT distribution has taken a relatively similar form to its fit on the full dataset by the time we have given it 25% of the data. The bottom row of Figure 1 shows the predicted frequency versus the observed frequency (see also Figure S1a for credible intervals based on posterior sampling). We see that the MAP estimate under SeqDEFT essentially matches the empirical frequency for states with greater than on the order of 10 observations, and thus the smoothness included by the prior essentially only influences our predictions in the more sparsely sampled regions of sequence space (as shown by deviations from the line *y* = *x*). With increasing data, the breakpoint between these two regimes moves to proportionally lower frequency states.

**Figure 1:**
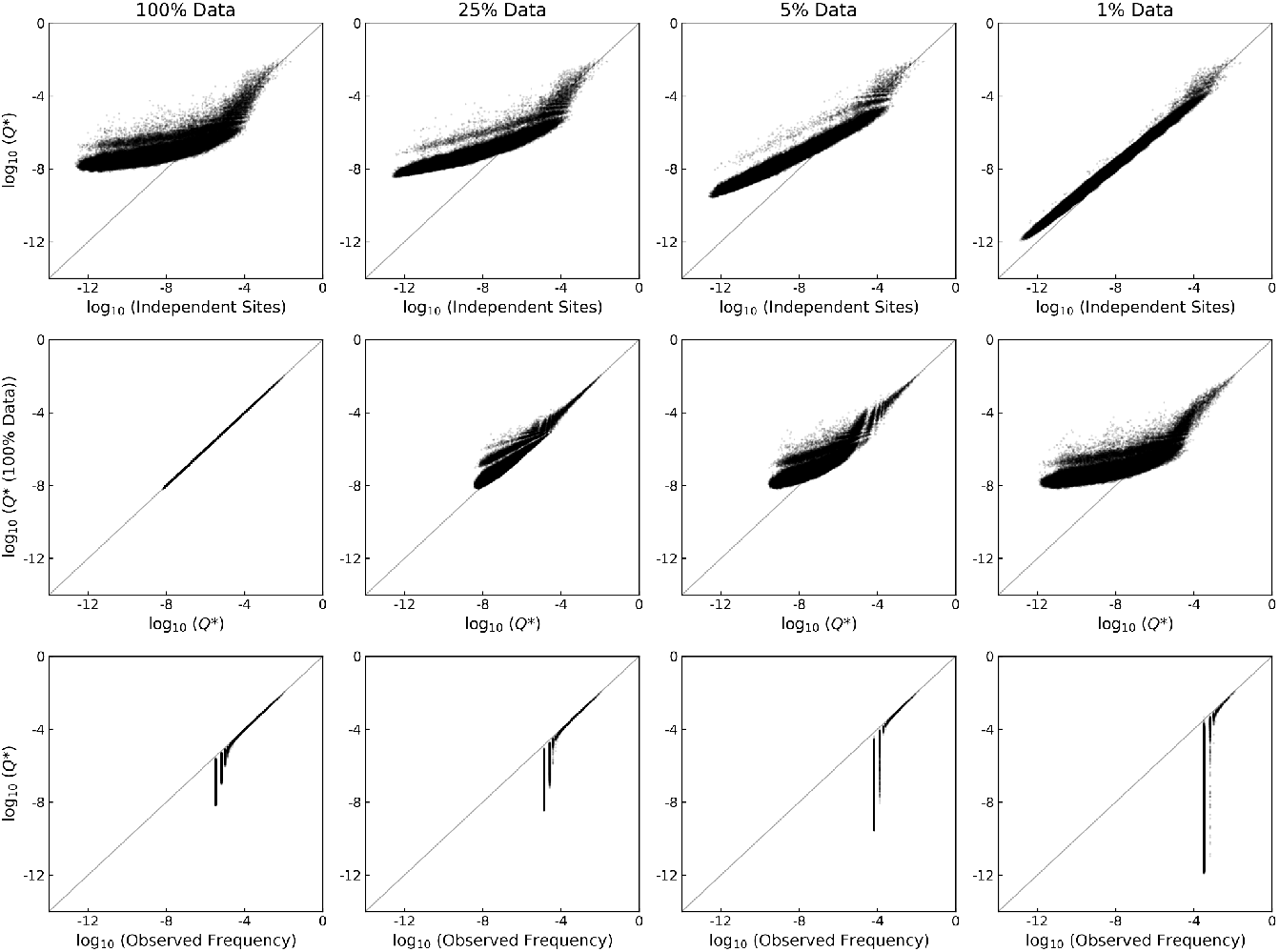
Behavior of SeqDEFT with changing sample size. The first row plots the SeqDEFT MAP estimate versus the independent positions maximum entropy model. The second row shows the SeqDEFT estimate using the full dataset versus the estimates for the smaller samples. The third row shows the SeqDEFT estimate versus the sampled frequencies. In order to show the class of sequences that were not observed in the sample we added a pseudo-count of 1 to the sequence counts. Thus the vertical striations indicated sequences that were observed 0 times, 1 times, 2 times, etc.@

To gain some intuition for this behavior it is helpful to consider the form of the posterior action. The first term 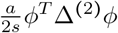 is the prior action and favors conditional bivariate distributions that are as nearly independent as possible. The second and third terms, 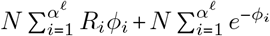, measure the match between *ϕ*_*i*_ and the observed data and are minimized by setting *ϕ*_*i*_ equal to the negative logarithm of the observed frequency *R*_*i*_. However, the value of these last two terms is relatively insensitive to the value of *ϕ*_*i*_ for *i* when the number of observations *NR*_*i*_ is zero, e.g. if *NR*_*i*_ = 0 then the corresponding *ϕ*_*i*_ does not contribute to the second term and contributes minimally to the third term as long as 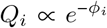 is small. Thus, the prior dominates in regions of sequence space where the number of observations is small, producing an MAP estimate with small conditional associations in these regions, whereas well-sampled genotypes are predicted to have frequencies similar to those that are empirically observed.

Another useful comparison is with the pairwise maximum entropy fit (Figure 2, left). We see that SeqDEFT and the pairwise maximum entropy model agree closely for high-frequency sequences, but that SeqDEFT produces a much narrower range of estimates for low-frequency sequences, and this latter pattern is in fact similar to what we observe for the independent sites maximum entropy model as well (Figure 1, top left).

**Figure 2:**
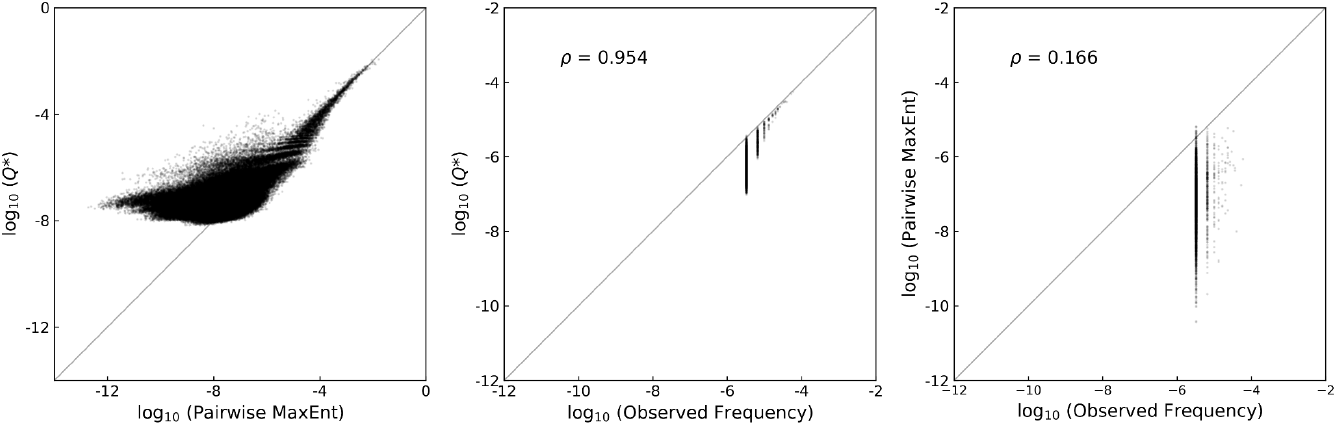
Comparison of SeqDEFT and pairwise maximum entropy fit to the distribution of human 5’ splice sites. The left panel shows the SeqDEFT MAP estimate versus the pairwise maximum entropy fit.The middle panel shows the SeqDEFT MAP estimate versus the empirically observed frequency for the subset of 5’ splice sites that do not have the canonical +1 G and +2 U/C nucleotides. The right panel shows the pairwise maximum entropy fit versus the observed frequency for the sub-set of 5’ splice sites that do not have the canonical +1 G and +2 U/C nucleotides.

Why do the maximum entropy models produce such a wide range of estimates in low-frequency regions of sequence space? The key observation is that for these maximum entropy models there is an assumption that some feature of the data remains constant over all of sequence space, e.g. for the independent model each possible mutation has a constant multiplicative effect on frequency and for the pairwise model the conditional log odds ratio between any given pair of mutations is constant over all of sequence space. Moreover, these constants are determined by the moments of the sample which are primarily influenced by the high-frequency sequences. Thus, if e.g. a pair of mutations are associated among the high-frequency sequences, the pairwise maximum entropy model will assume that they remain associated even in regions of sequence space where we have no data to support this association. In contrast, SeqDEFT allows these associations to decay in regions of sequence space where there is no data to support them, producing much smoother and more uniform predictions for poorly sampled regions. Such behavior is also in better accordance with biological intuition in that e.g. correlations between positions in functional sequences is due to natural selection on that function and thus these correlations should not be observed in regions of sequence-space that consist of non-functional sequences.

Another important difference between the maximum entropy models and SeqDEFT concerns SeqDEFT’s ability to learn components of the probability distribution that are at lower frequencies, e.g. in treating a multi-modal distribution, the maximum entropy solution will tend to fit the largest of the modes while providing a poor fit for other modes that might have strong statistical support but contain a small absolute fraction of the total probability. 5’ splice sites provide a good illustration of this principle. The vast majority of splice sites have a G in the +1 position and a U or C in the +2 position, but a small fraction (1.68% in our dataset) have other nucleotides [36, 37], and A in the +1 position in particular can be recognized by a different splicing machinery known as the minor spliceosome [38]. The center panel of Figure 2 shows SeqDEFT’s fit to these atypical, non +1 G or non +2 U/C, sequences while the right panel shows the fit of the pairwise maximum entropy model to these same sequences. We see that SeqDEFT is able to learn the density for these atypical sequences whereas the pairwise maximum entropy model produces a qualitatively incorrect fit.

An additional surprising feature of the SeqDEFT fit is that despite being more expressive it actually has smaller values for the inferred conditional log odds ratios than the pairwise maximum entropy model does. In particular, at the optimal value of *a* (in this case *a*∗ = 3.8 × 10^4^, Figure S2a) the MAP SeqDEFT model has a root-mean-square (RMS) log odds ratio of 0.656 and a cross-validated mean log likelihood of −424,970.8, whereas the pairwise maximum entropy model has an RMS log odds ratio of 0.864 and a cross-validated mean log likelihood of −429,167.8 (Figure S2b). The SeqDEFT fit is also slightly more concentrated in sequence space than the pairwise maximum entropy fit, a concept that can be quantified using the effective number of sequences, i.e. the number of sequences such that a uniform distribution over those sequences has the same entropy as our estimated entropy. In particular, posterior samples from SeqDEFT produce a 95% credible interval for the effective number of 5’ splice sites as being between 1019 and 1035, whereas the pairwise maximum entropy model corresponds to 1132 seqeuences and the independent sites maximum entropy model 1666 (see Table S1).

For pairwise maximum entropy models, in order to identify positions that may be interacting, it is common to construct heat maps showing the correlations between positions or the inferred log odds ratio for each pair of mutations. We can construct similar plots based on SeqDEFT estimates by displaying an appropriate summary statistic for the distribution of local interaction strengths, such as the RMS conditional log odds ratio for specific pairs of mutations. Figure 3 (left) shows a heat map of this type, and it is easy to see that the strongest interactions are between mutations altering the consensus +1 G and mutations altering the consensus G at the −1 position or the consensus U at +2 position, but it is also clear that there are interactions between many other pairs of mutations.

**Figure 3:**
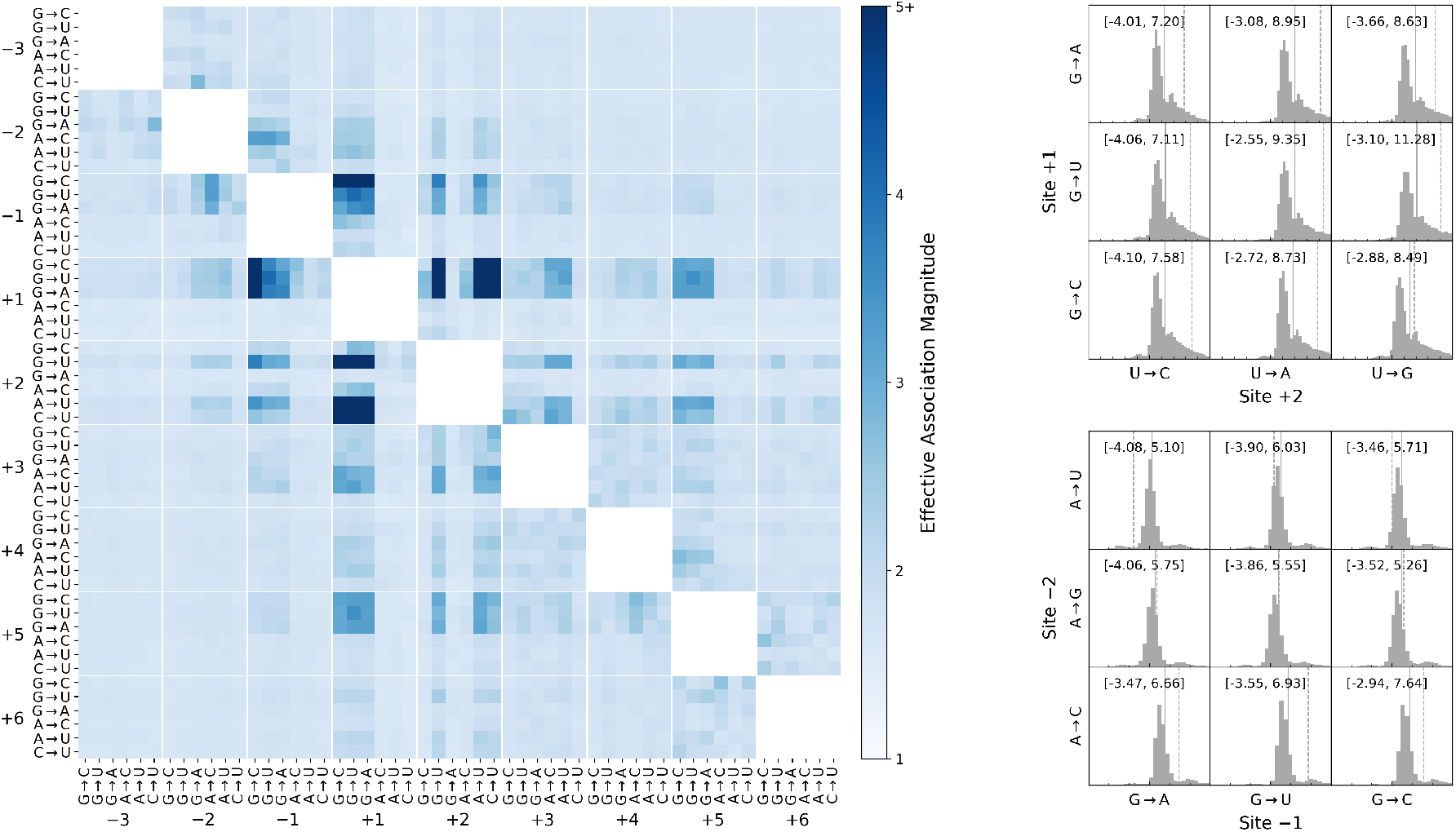
Interactions between pairs of mutations in the SeqDEFT fit. The left panel shows the exponentiated RMS log odds ratio between each possible pair of mutations under the MAP SeqDEFT fit. The RMS log odds ratio is calculated based on all faces of the Hamming graph corresponding to a given pair of mutations, and is exponentiated to express the effective strength of the association between any given pair of mutations on the same scale as the odds ratio. To provide a more fine-grained view of the variation in the strength of these association between different faces of the Hamming graph, the right panels show histograms of the distribution of odds ratios for mutations away from the consensus bases at the +1 and +2 positions (top-right) and −2 and −1 positions (bottom-right). The major tick on the x axis of each histogram indicates a log odds ratio of zero, and the other ticks are placed at intervals of 1 unit on the log-odds scale. To aid comparison, the histograms are shown with a limited range of log odds ratios on the x axis but the full range of inferred log odds ratios is shown by the bracketed values at the top of each panel. Mutations are polarized from the more preferred to the less preferred state, so that positive associations typically indicate that either of the single mutants results in loss of function while a negative association indicates that the two mutations are tolerable individually but not jointly. The solid vertical lines indicate the mean of the log odds ratios while the dotted vertical lines indicate the uniform log odds ratio assigned to that pair of mutations under the pairwise maximum entropy model.

However, unlike the pairwise maximum entropy model which must produce a single conditional log odds ratio for any given pair of mutants, SeqDEFT allows the strength and direction of the association between each pair of mutations to vary with the genetic background. Thus, rather than look at single summary statistic of association strength between pairs of mutations, we can also take a more detailed view by plotting histograms of the log odds ratios inferred for different genetic backgrounds (i.e. how the log odds ratio varies over the different faces of the Hamming graph). Figure 3 right-top shows these distributions for mutations to the consensus nucleotides at the +1 and +2 positions. We see that these histograms are strongly right-skewed, a pattern that likely arises because mutations to either of these nucleotides typically renders a functional splice site non-functional (producing a strong positive association in the conditional distribution) but has little effect on an already non-functional splice site. The solid vertical line in the plots indicates the mean conditional log odds ratio (averaged over all faces in the Hamming graph) while the dashed vertical line indicates the constant log odds ratio for all such faces assigned by the pairwise maximum entropy model. Figure 3 right-bottom shows a similar set of distributions for interactions between the −1 and −2 positions. A careful examination of these histograms shows that they are typically tri-modal, with a large central mode and two smaller side modes, indicating that a subset of sites show a substantial interaction between these two mutations, but that the sign of this association differs depending on the genetic background. Thus, our SeqDEFT estimate suggests that the sign and strength of the association between a pair of mutations can vary in a complex manner depending on the genetic background, an observation that is qualitatively incompatible with the assumptions of a pairwise maximum entropy model.

### Visualizing the inferred geometry using an evolutionary model

A key strength of non-parametric approaches such as SeqDEFT is that, provided sufficient data, they can capture whatever complex geometry is present in the data. However, this comes at the expense of interpretability, because we can no longer express the inferred distribution in terms of a small number of parameters. We have already explored one way of overcoming this difficulty, in the form of the summary statistics and histograms shown in Figure 3. A different solution is to attempt to visualize or represent the inferred distribution in such a way that these visualizations allow us to identify the major qualitative features of the distribution and explore the underlying causes of these features.

The visualization approach we take here [39] is based on considering our inferred probability distributions over sequence space as being the result of an evolutionary process (SI Appendix for details). The main idea is that biological evolution can be viewed as the process of a population taking a random walk over sequence space, where each step in the random walk consists in the replacement of one sequence in the population by another, and the role of natural selection is to bias the probability that any given mutational neighbor of the current sequence becomes fixed [40, 41]. With this idea in mind, given an inferred distribution over sequence space we can write down a model of molecular evolution as a Markov chain which takes that distribution as its stationary distribution. To do this, we note that in certain standard models of molecular evolution the quantity log 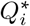 is equal to the product of the fitness of state *i* and the effective population size, so that population genetic theory predicts that the transition rate between *i* and *j* is given by

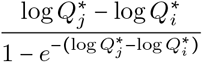

for mutationally adjacent sequences *i* and *j* [40, 42]. Using these transition rates, we can then construct visualizations of the resulting evolutionary dynamics by plotting each genotype at coordinates given by certain subdominant eigenvectors of the rate matrix of the Markov chain, and we refer to these axes as diffusion axes [43] because they capture the slow modes of the diffusion of the probability distribution describing the location of the population in sequence space. Moreover, by scaling these axes appropriately, the squared Euclidean distance between genotypes *i* and *j* optimally represents the expected time to evolve from one genotype to the other so that the axes in the visualizations have units of square-root time (here, we set the units of time so that each possible point mutation occurs at rate 1).

In the case at hand, we are considering a population evolving under selection to have a functional splice site at a particular location in the genome under the assumption that the stationary distribution for this process is given by the inferred distribution of 5’ splice site sequences given by SeqDEFT. Figure 4 shows the resulting visualization, where we have fixed the +1 and +2 positions to be the canonical GU nucleotides in order to best display the geometrical features of the subset of functional sequences (see Supplemental Figure S3 for the corresponding visualizations over all of sequence space).

**Figure 4:**
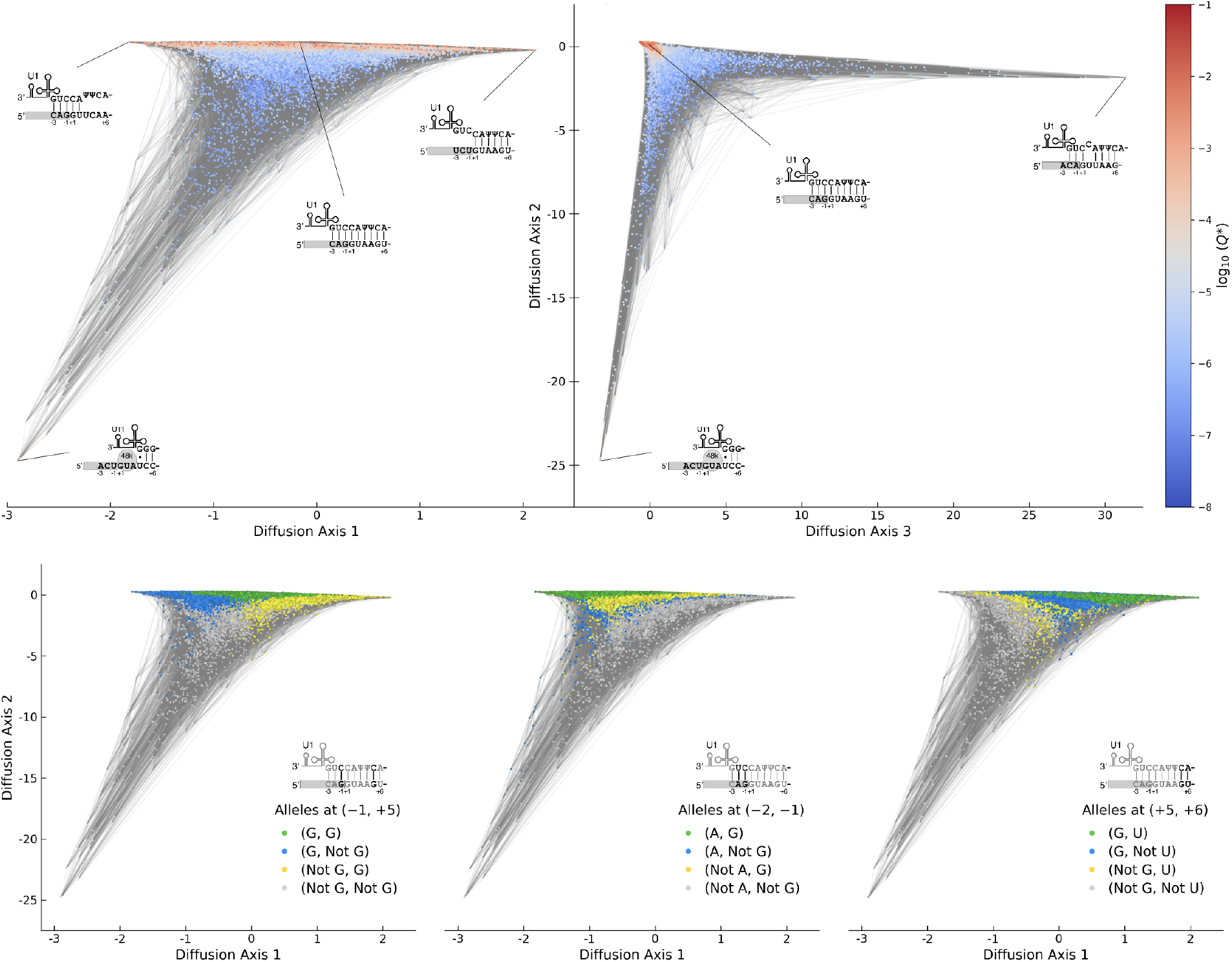
Visualization of the distribution of 5’ splice sites inferred by SeqDEFT. Visualization uses the method of [39] where the log 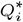 is equated with the scaled fitness of genotype *i*, and this quantity is used to define an evolutionary Markov chain whose stationary distribution corresponds to the MAP SeqDEFT estimate (conditioned on the +1 and +2 positions being the canonical GU nucleotides). Sequences are colored according to their estimated frequency and edges connect sequences that are adjacent under point mutations. Under the visualization method squared Euclidean distances optimally approximate the expected time to evolve from one sequence to another and we scale time so that each possible point mutation occurs at rate 1. The top row shows the first 3 diffusion axes, which are re-scaled subdominant eigenvalues of the transition matrix for the Markov chain. The cartoons show hypothesized binding geometries. In the bottom row, sequences are colored to show the base identities at the positions shown in black in the corresponding cartoon. See Supplemental Figure S5 for the corresponding visualization based on the pairwise maximum entropy density estimate, which does not show any obvious structure.

The top row of Figure 4 shows the first 3 diffusion axes. Genotypes are colored according to their inferred frequency and edges connect genotypes that differ by a single point mutation. We see that there is a connected cluster of sequences similar to the canonical binding motif CAG/GUAAGU that is stretched along Diffusion Axis 1, and then two clusters of moderate frequency with extreme values on Diffusion Axis 2 or Diffusion Axis 3. To understand the structure of the main streak, it is helpful to know that the canonical 5’ splice site is recognized and bound by a small RNA known as the U1 snRNA with which it forms a series of adjacent basepairs resulting in a helical structure [35]. Moreover, it has been previously observed that 5’ splice sites show a pattern of 5’/3’ compensation or “seesaw” linkage where mismatches in the exonic portion of the binding site are associated with consensus nucleotides in the intronic portion and vice versa [44, 45, 46]. The long axis of the streak turns out to correspond to this pattern in the locations of mismatches between the 5’ splice site and U1, with sequences that primarily form basepairs in the exonic portion of the splice site having negative values on Diffusion Axis 1 and sequences that bind via basepairs in the intronic portion having positive values (see Supplemental Figure S4 for the average position of consensus bases for sequences in the streak as a function of position along Diffusion Axis 1). The bottom panels show this geometry in more detail, demonstrating that the streak of high frequency sequences have at most one mismatch at either the −1 or +5 position, and that the sequences with the most negative values on Diffusion Axis 1 have the canonical matching alleles at positions −2 and −1 while the sequences with positive values for Diffusion Axis 1 have the correct nucleotides at positions +5 and +6. These two sides of the streak are broadly separated because it would take many mutations to transform a sequence with primarily exonic basepairs into a sequence with primarily intronic basepairs and mismatches in the exonic portion of the binding site.

Besides the main streak of sequences that appear to bind via minor variations on the canonical 5’ splice site motif, there are two smaller clusters of high frequency sequences plotted far from the main streak. The cluster with a negative values on Diffusion Axis 2 corresponds to sequences that are recognized by the minor spliceosome [38]. This machinery recognizes the 5’ splice site via binding with the U11 rather than U1 snRNA, and this recognition occurs in conjunction with another protein known as 48K where the 48K protein contacts the +1 and +2 positions and binding with U11 is completely intronic, beginning at the +3 position [47]. The other small cluster of high frequency sequences turns out to correspond to a known non-canonical binding geometry known as the shifted +1 register [48] where U1 snRNA binding forms a gapped helical structure that is shifted one basepair over from the canonical motif, but where the position of the splicing reaction (transesterification) itself remains unchanged.

Now that we have identified the major geometric features of the inferred probability distribution and identified each of them as corresponding to a distinct biophyiscal mechanism of splice site recognition, we can also use these visualizations to ask questions about the evolution of binding by considering different paths that populations can take through sequence space. For example, it has been previously observed in genomic comparisons between different species that individual splice sites can be “converted” from being recognized by the minor spliceosome to being recognized by the major spliceosome and vice versa [49, 50]. In our visualizations we see that the sequences recognized by the minor spliceosome have large negative values on Diffusion Axis 1 and that for sequences similar to the canonical motif, Diffusion Axis 1 separate sequences where U1 binds primarily to the exonic portion of the splice site (negative values on Diffusion Axis 1) from those where it binds primarily in the intronic portion (positive values on Diffusion Axis 1). This suggests that conversion is most likely to occur via a transition from minor spliceosome recognition to a sequence capable of being recognized by U1 via the gain or loss of a CAG motif in positions −3 to −1 (i.e. paths leading up or down the left side of Figure 4). Similarly, we can ask about the even longer time possibility of transitions between all three clusters. An important characteristic of these visualizations is that because the visualizations attempt to capture the expected time it takes to evolve from one genotype to another by the squared Euclidean distance, chains of sequences A-B-C where A typically evolves to C via B intermediates appear as right angles (since the time to evolve from A to C will approximately equal the sum of the time to evolve from A to B plus the time to evolve from B to C). Thus, the right angle pattern between the three clusters in the right panel of Figure 4 suggests that transitions between the minor spliceosome and the shifted +1 U1 binding register will tend to occur via evolutionary intermediates that bind U1 in the canonical register rather than by direct transitions between minor spliceosome and +1 shifted U1 binding.

### Distribution of karyotypic abnormalities in cancer

Our example of human 5’ splice sites provides both a demonstration of SeqDEFT’s behavior in the well-sampled regime and of the rich geometry of biological distributions that can be captured via a non-parametric approach. We now turn to an example in the poorly-sampled regime, where the number of sequences far exceeds the number of observations. In particular, we consider the problem of karyotypic abnormalities in human cancers, where cancerous cells frequently exhibit a bewildering array of changes to the structure and number of chromosomes ranging from losses and duplications of small portions of individual chromosomes, to duplications and losses of chromosome arms, translocations that attach a portion of a chromosome to another, duplications and losses of whole chromosomes, and (multiple) whole genome duplications [51]. Moreover, the root causes of these changes in genomic structure remain poorly understood because chromosomal copy number changes can either promote or inhibit cellular proliferation depending on the specific alterations involved [52, 53, 54].

To better understand the distribution of karyotypic states exhibited by human cancers, we considered karyotypes inferred for 10,522 tumors [29] collected as part of The Cancer Genome Atlas [55]. Starting with the simplest possible approach, we considered each of the 22 autosomes and scored each autosome as being either euploid or aneuploid, where we scored a chromosome as aneuploid if [29] reported that chromosome as exhibiting large-scale alterations from the background cellular ploidy. This scoring scheme results in a total of 2^22^ = 4,194,304 possible karyotypic states.

In fitting this data with SeqDEFT, we observed that the pairwise maximum entropy model had a greater cross-validated log likelihood than our model using Δ^(2)^ (i.e. the pairwise maximum entropy model provided a higher likelihood fit than a model based on a perturbation of the independent model, SI Figure S2c,d). This was likely caused by a particularly strong global pattern of non-independence between chromosomal states wherein observed karyotypes were roughly uniformly (rather than binomially) distributed in terms of their number of aneuploid chromosomes (SI Figure S6a), indicating a strong enrichment for chromosomal configurations that are either perturbations of the standard euploid genome with a handful of altered chromosomes or else nearly completely aneuploid (SI Figure S6b). This observation is consistent with the well-known phenomenon of chromosomal instability, where deviations from a euploid karyotype result in further mitotic errors and hence an increasingly high degree of aneuploidy [56]. We therefore proceeded with an analysis based on Δ^(3)^, which relaxes the constraints of the pairwise model. Importantly, the pairwise maximum entropy model treated all chromosomes in an approximately uniform manner, where the marginal frequencies of aneuploidy for each chromosome have a mean of 0.41 and standard deviation of 0.06 for each chromosome and conditional log odds ratios with a mean of 0.23 and a standard deviation of 0.17, so that the presence of any one chromosomal alteration increases the probability of each of the others by an approximately constant factor. Thus, while the maximum entropy model does a good job capturing the overall bimodality of the data, it does not appear to be capturing more detailed interactions between specific pairs or subsets of chromosomes.

Using Δ ^(3)^ we find that *a*^*∗*^ *=* 7.9 × 10^5^, resulting in an increase of 422.4 in the cross-validated log likelihood relative to the pairwise maximum entropy model and a moderate decrease in the estimated effective number of cancer karyotypes, from 1.1 × 10^6^ for the maximum entropy model to a 95% credible interval of 5.9 × 10^4^ to 7.2 × 10^4^ based on posterior sampling (Table S1, see Supplemental Figure S1b for posterior variance estimates). Although the SeqDEFT predictions are mostly similar to the pairwise maximum entropy model there are also relatively dramatic differences for a subset of sequences that SeqDEFT predicts to be at much higher frequencies than the pairwise maximum entropy model (SI Figure S6c). To better understand what these high frequency sequences are, we turn to our visualization technique [39]. Here our application of the visualization technique is somewhat less principled since a cancer’s exploration of sequence space is not stationary but rather ends in either the patient’s death or eradication of the tumor, and there are a number of other differences such as e.g. gains or losses of multiple chromosomes at once (so that evolution is not restricted to the edges of the Hamming graph) and polyclonality within the tumor (so that the tumor contains cells with a number of different karyotypic states rather than a single state for each time). Nonetheless, the visualizations do provide insight into the basic geometry of the inferred probability distribution and hence indirectly into the process that generated it.

Figure 5 (top row) shows the resulting visualization. Here, Diffusion Axis 1 picks out the number of chromosomes that are altered with the wild-type euploid karyotype on the left and the karyotype where all chromosomes are aneuploid on the right (faint striations are also visible which correspond to Hamming distance from the wild-type sequence). However Diffusion Axis 2 reveals two regions of unusually high frequency sequences in a region of sequence space where most sequences are much lower frequency. In particular, the tip sequences for these protrusions are karyotypes with simultaneous copy number changes at chromosomes 1, 2, 6, 10, 13, 17, and 21 or at chromosomes 6, 7, 9, 10, 19, and 20. Diffusion axis 3 then reveals the geometric relationship between these high frequency regions, showing that they are two distinct regions that both branch off the main arc that connects the wild-type to the all aneuploid state. The top right panel of Figure 5 also shows a protrusion of high frequency sequences around the state with chromosomes 2, 3, 7, 12, 16, 17, 20, and 21 simultaneously altered, which turns out to appear as a third branch-like protrusion when we include Diffusion axes 4 and 5 in our visualizations (SI Figure S7).

**Figure 5:**
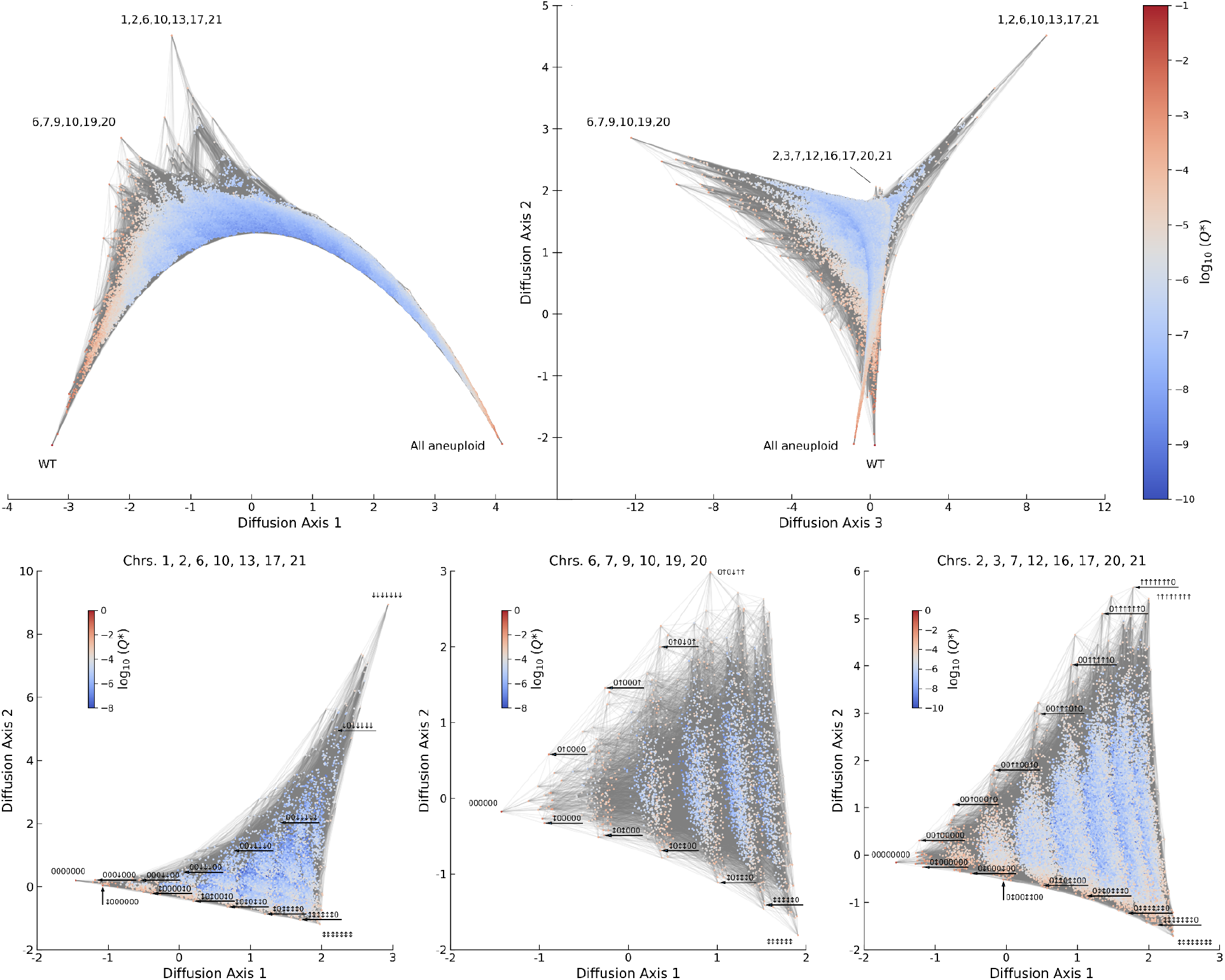
Visualization of the distribution of human karyotypes based on data from [29]. Visualization uses the method of [39] where the log 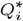 is equated with the scaled fitness of karyotype *i*, and this quantity is used to define an evolutionary Markov chain whose stationary distribution corresponds to the MAP SeqDEFT estimate. Sequences are colored according to their estimated frequency and edges connect sequences that can be transformed one into the other by changing the state of a single chromosome. The top row shows the first 3 diffusion axes for a binary encoding of the karyotypic state where each of the 22 autosomes is scored as being euploid or aneuploid; the pattern of aneuploid chromosomes is indicated for key local maxima in the estimated probability distribution. See SI Figure S7 for Diffusion axes 4 and 5. The bottom row shows visualizations based on *Q^*^* when estimated using data from only the indicated chromosomes which are encoded in a 4-state scheme, with states gain (up arrow), loss (down arrow), complex change (double-headed arrow) or euploid (0). Genotypes are labeled along the paths from the all euploid state to either the focal pattern of aneuploidy or the all complex aneuploid genotype, where the selected path is the minimal length path defined by moving to the highest frequency neighbor at each step.

What do these visualization tell us about the geometry of our inferred probability distribution? Our evolutionary Markov chain treats log frequency as a measure of evolutionary fitness. Thus, it treats the wild-type and all aneuploid sequences as two major fitness peaks, separated by the broad valley of partially aneuploid sequences, so that populations typically stay at one of these two peaks or the other, but occasionally stochastically transition from one to the other. However, within this valley, we have observed three clusters of high frequency sequences, with local frequency (and hence fitness) maxima at states with chromosomes {1, 2, 6, 10, 13, 17, 21} or {6, 7, 9, 10, 19, 20} or {2, 3, 7, 12, 16, 17, 20, 21} altered. Populations that wander into the basin of attraction of these local fitness maxima can become stuck there, leading to long waiting times for these populations to visit the other maxima and hence large distances between these maxima in our visualizations. Importantly, using the pairwise maximum density estimate for the visualization does not reveal any of this fine-scale structure (SI Figure S8), which demonstrates the power of non-parametric approaches to reflect the complexities of biological data.

To better understand the biological basis of these local maxima in our inferred probability distribution, we also considered which specific tissue types contributed to these maxima. We found that the regions around {1, 2, 6, 10, 13, 17, 21}and {2, 3, 7, 12, 16, 17, 20, 21} were composed of cancers that originated in kidney tissue and that the region around {6, 7, 9, 10, 19, 20} was based in observations of brain cancers, with other tissue types making little or no contribution to these local maxima (SI Figure S9). Furthermore, we found that the simultaneous chromosomal alterations at chromosomes {1, 2, 6, 10, 13, 17, 21} or {2, 3, 7, 12, 16, 17, 20, 21} were in fact largely due to different kidney cancers, with the enriched frequency around {1, 2, 6, 10, 13, 17, 21} primarily due to chromophobe renal cell carcinoma (KICH) and the enriched frequency around {2, 3, 7, 12, 16, 17, 20, 21}primarily due to kidney renal papillary cell carncinoma (KIRP), with kidney clear cell carcinoma (KIRC) making some contribution to both regions (SI Figure S10). At the same time, we see that for both brain and kidney tissues many tumors also exhibit much more extensive aneuploidy, suggesting a dynamic where individual tumors can become attracted towards certain specific combinations of chromosomal abnormalities before ultimately progressing to a more highly aneuploid state.

Another important question is the specific set of chromosomal abnormalities that contribute to each of these high frequency signatures, since our simple binary scoring lumps together chromosomal losses and gains. Increasing to even three states (euploid, gain, loss) for all 22 autosomes would result in a sequence space containing more than 31 billion sequences, which is too large for our current sparse matrix multiplication-based computational implementation (see Supplemental Figure S11 for qualitatively similar results obtained under 2 alternative binary scoring schemes). However, we can consider a larger number of states per chromosome while restricting the genotype to consist of only the specific subsets of chromosomes that appear to be interacting based on our above analysis. In particular, Taylor et al. [29] provide calls for each chromosome as being euploid or having an increase, decrease, or complex aneuploidy (e.g. gain of one chromosome arm but not the other or a more complex mixture of amplified and deleted chromosomal segments), and we re-ran our SeqDEFT analysis with this new scoring system for each of the three focal combinations of chromosomes identified above.

The qualitative form of each of these inferred probability distributions is similar, as shown using the visualizations in the bottom row of Figure 5. In each case, the first diffusion axis captures the total number of aneuploid chromosomes in a genotype, and then the second axis distinguishes two different forms of accumulation of aneuploid chromosomes: one consisting of complex aneuploidy for all chromosomes (with complex aneuploidy indicated by a double-headed arrow pointed both up and down) and the other consisting of a specific pattern of gains (up arrows) or losses (down arrows). For {1, 2, 6, 10, 13, 17, 21}, the specific attracting pattern appears to be losses of all 7 of these chromosomes, a pattern that has previously been recognized as a signature of chromophobe renal cell carcinoma [57]. For {6, 7, 9, 10, 19, 20}, the pattern is more complicated with the main attracting state being gains of chromosomes 7, 19 and 20 together with loss of chromosome 10, where simultaneous gains of chromosomes 7 and 19 together with loss of chromosome 10 is widely known to be common in glioblastomas [58] and co-amplification of chromosomes 19 and 20 has been identified as a marker of positive prognosis [59]. Then this pattern at chromosomes 7, 10, 19 and 20 is frequently complemented, particularly in glioblastomas, by either loss or complex aneuploidy of chromosome 9, and then in the presence of this additional chromosomal change we also frequently see loss or complex aneuploidy of chromosome 6 in glioblastoma. So in this case the “peak” sequence in the binary encoding appears to correspond to a cloud of several different karyotypes in our more detailed encoding, where sequences in this cloud all share the same core motif but with different elaborations. For {2, 3, 7, 12, 16, 17, 20, 21}, the observed pattern is simultaneous copy number gains for all these chromosomes, a pattern that has also been noted in the literature as being a signature of renal papillary cell carcinoma [60].

Besides identifying higher-order combinations of chromosomal abnormalities that occur at high frequencies, we can also ask questions about how these configurations evolve from the initial euploid state. The visualizations in the bottom row of Figure 5 show some examples of likely high probability trajectories; in particular, for each combination of chromosomes we have labeled the minimal length path to the signature pattern of gains and losses and the minimal length path to the genotype where all chromosomes are altered under the assumption that the cancer population always incorporates whichever of the remaining mutations would result in the highest frequency genotype.

To summarize, even with relatively few observations compared to the overall size of sequence space for karyotypic abnormalities, we are able to identify specific patterns of chromosomal gains and losses that appear to be favored in specific tissues, and we can begin to ask questions about the geometric relationships between these high frequency regions and the order in which the constituent mutations accumulate during cancer progression.

## Discussion

Probability distributions in molecular biology are often complex and idiosyncratic because they inherit the complexity and idiosyncrasy of the chemical, historical, and evolutionary processes from whose confluence they arise. Likewise, the character of these probability distributions is often discrete and combinatorial because the organization of biological information typically takes this form, either in the guise of informational heteropolymers (RNA, DNA, proteins) or because biological complexity often takes the form of a collection of subunits where each subunit can be in a certain number of states (a collection of neurons, each of which is firing or not; a collection of chromosomes that each appears with a certain number of copies). Here we have proposed a flexible method for estimating probability distributions over these types of discrete combinatorial spaces that is capable of capturing the detailed idiosyncrasy typical of these naturally occurring distributions. If we consider the simplest and most familiar probability distributions such as exponential and normal distributions, these distributions often arise as maximum entropy distributions, that is the most uniform distributions compatible with some set of constraints on the moments of the distribution (e.g. the normal distribution is the most uniform distribution given a fixed mean and variance). Recent advances in Bayesian field theoretic density estimation [22, 23, 24, 25, 26, 27, 28] have shown that it is possible to elaborate on such maximum entropy estimates by defining a suitable prior over the space of possible probability distributions. The theory we have developed here has a completely analogous structure, but transferred to a discrete multivariate setting (SI Table S2). For example, in the continuous case, the key quantity for determining the prior probability of a particular probability distribution is the integral of the squared *P* −th order derivative of the log density, which is a measure of the average local roughness of the probability distribution. Here the corresponding quantity is e.g. the average squared value of the conditional log odds ratio. This makes perfect sense once one realizes that the conditional log odds ratio is just the discrete analog of a second order mixed partial derivative, since it measures (locally) how changing the value of one covariate alters the effect of changing another covariate.

The work proposed here is also closely related to Minimum Epistasis Interpolation, a technique that we recently proposed for regression, rather than density estimation, in biological sequence space [30]. The relationship between these techniques is that if we view − *ϕ*_*i*_ as a phenotype, then the conditional log odds ratio exactly corresponds to the classical notion of a double mutant epistatic coefficient. Minimum Epistasis Interpolation works to estimates the relationship between genotype and phenotype by taking a set of known phenotypic values and estimating the remaining values by minimizing the value of the average squared double mutant epistatic coefficient, i.e. minimizing *ϕ*^*T*^ Δ ^(2)^ *ϕ* subject to an equality constraint at known genotypes. This can produce a very complex reconstruction in regions of sequence space where data is plentiful, but relaxes towards a non-epistatic (i.e. additive) reconstruction in regions of sequence space where data is sparse or absent. In the regression case, the solution to this problem is given by solving a single system of equations, unlike the nonlinear problem explored here. However, at an intuitive level, the SeqDEFT problem is very similar to Minimum Epistasis Interpolation in that sequences with large number of observations have essentially known log frequencies whereas sequences with zero or low counts have essentially unknown log frequencies, and the prior works to ensure that the model displays relatively simple behavior in these poorly sampled regions while matching our observations in well-sampled regions.

Our results here also provide some immediate extensions to our previous results on Minimum Epistasis Interpolation. First, our results provide an intuitive perspective on the Bayesian generalization of Minimum Epistasis Interpolation by demonstrating that its prior is equivalent to drawing epistatic coefficients independently for each face of the Hamming graph and then conditioning on mutual consistency (i.e. being a valid solution to the bookkeeping problem, [31]). Second, the Δ ^(*P*)^ for *P >* 2 can be used immediately to provide higher-order analogs of the original Minimum Epistasis Interpolation technique by calculating the constrained minimizer of *ϕ*^*T*^ Δ ^(*P*)^ *ϕ*.

Although SeqDEFT can capture complex probability distributions in the bulk of the data while exhibiting simpler behavior in poorly sampled regions of sequence space, this flexibility comes at a cost in terms of computational complexity. The main issue is the non-quadratic nature of the posterior action, which results in a numerical minimization problem that in practice we can only currently solve for sequence spaces with a few million genotypes or less. This limitation corresponds to a maximum length of 11 for DNA sequences or 5 for amino acid sequences, far from the whole-protein scale where pairwise maximum entropy models have shown such impressive performance [14, 19, 20, 21]. More work is needed to develop non-parametric inference and exploratory data analysis techniques applicable to these larger sequence spaces.

## Materials and Methods

Our implementation of SeqDEFT is available at https://github.com/davidmccandlish/SeqDEFT. Sparse matrices and their manipulations were computed using the SciPy “sparse” package. MAP solutions were estimated by minimizing the posterior action using the SciPy “optimize” package. The optimal hyper-parameter was determined by maximizing the *k*-fold cross-validated log likelihood with *k* = 5. Probability distributions were visualized using the dimensionality reduction technique in ref. [39]. Details are given in SI Text.

The dataset of annotated human 5’ splice sites was extracted from GENCODE Release 34 (GRCh38.p13), available at https://www.gencodegenes.org/human/. The dataset of karyotypic abnormalities in human cancer was from ref. [29] and is available as part of the supplementary material at https://doi.org/10.1016/j.ccell.2018.03.007.

## Acknowledgments

This work was supported by NIH Cancer Center Support Grant No. 5P30CA045508, NIH Grants R35GM133613 (D.M.M.) and R35GM133777 (J.B.K.), an Alfred P. Sloan Research Fellowship (D.M.M.), a CSHL/Northwell Health Alliance grant (J.B.K.), and also by additional funding from The Simons Center for Quantitative Biology at Cold Spring Harbor Laboratory.

