## Supplemental Information for "Non-parametric Bayesian density estimation for biological sequence space with applications to pre-mRNA splicing and the karyotypic diversity of human cancer"

#### Formalism of SeqDEFT

Here we provide a self-contained description of our approach, Sequence Density Estimation using Field Theory (SeqDEFT). This approach is a combination of Bayesian field theory and spectral graph theory. Detailed derivations of quantities related to Bayesian field theory can be found in [1]. A thorough introduction of the graph theory techniques employed in this work can be found in [2]. Given that this work can be regarded as an extension of the field-theoretic one-dimensional density estimation, we provide a heuristic comparison between these two methods in Table 2.

##### Setup

Consider sequences of length $\ell$, with each site of the sequences having $\alpha$ alleles. The number of all possible sequences is $G = \alpha^\ell$. These sequences together form a discrete combinatorial space. Given a set of count data of the sequences, our goal is to infer the underlying probability distribution from which the observed sequences were drawn.

We can represent the sequence space as a Hamming graph formed by a $\ell$-fold Cartesian product of the $\alpha$ alleles. Thus each sequence, represented as a node in the graph, has a Hamming distance 1 to all its neighbors. Such a graph has some intriguing properties. In particular, its Laplacian matrix $L$ (with $L_{ij} = -1$ if $i$ and $j$ are mutational neighbors, $\ell(\alpha-1)$ if $i = j$, and zero otherwise) has the following analytic spectrum:

$$L u_k = \lambda_k u_k, \qquad \lambda_k = k\alpha, \qquad m_k = \binom{\ell}{k}(\alpha-1)^k \tag{1}$$

for $k = 0, 1, 2, \ldots, \ell$, where $\lambda_k$ is the $k$th eigenvalue of $L$ with multiplicity $m_k$ and $u_k$ is the corresponding eigenvector. The Laplacian matrix, or graph Laplacian, provides a measure of "smoothness" of functions defined on the graph. Specifically, evaluating the quadratic form defined by $L$ on a function $\phi$ on the graph results in

$$\phi^T L\, \phi \equiv \sum_{i,j=1}^{G} \phi_i L_{ij} \phi_j = \sum_{e_{ij}} (\phi_i - \phi_j)^2 \tag{2}$$

where $e_{ij}$ is the edge connecting node $i$ and node $j$, and each edge is counted only once. A large value of $\phi^T L\, \phi$ means that $\phi$ is rather rough on the graph, whereas a small value of $\phi^T L\, \phi$ indicates that $\phi$ is close to uniform.

*

The edge $e_{ij}$ in Eq. 2 corresponds to certain mutation at a particular site of some specfic sequence. This geometric notion of edge can be generalized to higher-dimensional "hypercubes" (e.g., an edge can be viewed as a one-dimensional hypercube). On the other hand, the fact that the Laplacian matrix $L$ of the graph representing the sequence space of interest has a simple analytic spectrum allows us to define a series of matrices (or operators) of different order $P$, $P = 1, 2, \ldots, \ell$, in the following manner:

$$L^{(P)} = (L - \lambda_0)(L - \lambda_1)(L - \lambda_2)\cdots(L - \lambda_{P-1})\,. \tag{3}$$

The eigenvalue of the matrix $L^{(P)}$, associated with eigenvector $u_k$, is given by

$$\Lambda_k^{(P)} = (\lambda_k - \lambda_0)(\lambda_k - \lambda_1)(\lambda_k - \lambda_2)\cdots(\lambda_k - \lambda_{P-1}) \tag{4}$$

for $k = 0, 1, 2, \ldots, \ell$. It is clear that $\Lambda_k^{(P)} = 0$ for $0 \le k \le P - 1$. Thus, the kernel of $L^{(P)}$ enlarges as $P$ is increased. These generalizations, namely, the notion of hypercubes and the introduction of $L^{(P)}$, enable us to generalize Eq. 2 to compute higher-order epistatic coefficients (where we are viewing $\phi$ as a phenotype defined over sequence space). We outline this result in the following and give the proof in the Appendix.

Denote the $P$-dimensional hypercube by $P$-cube, $P = 1, 2, \ldots, \ell$. Let $m = \{m_1, m_2, \ldots, m_P\} \subset \{1, 2, \ldots, \ell\}$ be a subset of $P$ mutation sites, $A = \{A_{m_1}, A_{m_2}, \ldots, A_{m_P}\}$ a set of subsets with $A_{m_k} \subset \{1, 2, \ldots, \alpha\}$ and $|A_{m_k}| = 2$ for $k = 1, 2, \ldots, P$, and $b$ the "background" sequence of length $(\ell - P)$. The set $A$ provides a choice of the two alleles at each of the $P$ mutation sites. Further, let $c_{b,A}^m$ denote the $P$-cube consisting of the $2^P$ genotypes specified by $m$, $A$, and $b$. This $P$-cube can be formed by a Cartesian product, that is, $c_{b,A}^m = A_{m_1} \times A_{m_2} \times \cdots \times A_{m_P} \times \{b_1\} \times \{b_2\} \times \cdots \times \{b_{\ell-P}\}$, and the total number of such $P$-cubes is given by

$$s = \binom{\ell}{P}\binom{\alpha}{2}^P \alpha^{\ell-P}. \tag{5}$$

Within each hypercube, we have a bi-allelic sequence space of length $P$. Therefore, we can encode a sequence $\sigma$ of the hypercube by a binary sequence $\sigma'$ of length $P$, where the assignment of the two alleles in $A_{m_k}$ to 0 or 1 can be done arbitrarily. Define the $P$th-order epistatic coefficient of a function $\phi$ over the hypercube $c_{b,A}^m$ as

$$\epsilon_{c_{b,A}^m}(\phi) = \sum_{\sigma \in c_{b,A}^m} (-1)^{h(\sigma')}\phi(\sigma) \tag{6}$$

where $h(\sigma')$ gives the Hamming weight (i.e., the number of nonzero sites) of the sequence $\sigma'$. The mean squared $P$th-order epistatic coefficient of the function is then equal to

$$\epsilon_P^2(\phi) = \sum_m \sum_A \sum_b \epsilon_{c_{b,A}^m}^2(\phi)\,. \tag{7}$$

Notice that $\epsilon_P^2(\phi)$ consists of weighted sums of terms quadratic in $\phi$. This leads us to the following result:

**Proposition 1.** *$\epsilon_P^2(\phi)$ defines a positive semi-definite quadratic form which can be written as the following:*

$$\epsilon_P^2(\phi) = \phi^T \Delta^{(P)} \phi \tag{8}$$

*where $\Delta^{(P)}$ can be found to be*

$$\Delta^{(P)} = \frac{1}{P!} L^{(P)} = \frac{1}{P!}(L - \lambda_0)(L - \lambda_1)(L - \lambda_2)\cdots(L - \lambda_{P-1})\,. \tag{9}$$

Notice that while in the main text $\Delta^{(P)}$ is given an explicit definition from the outset, here we show that the formula presented in the main text can in fact be derived from the definition of the $P$-th order epistatic coefficient. The proof is provided as Appendix 1 below but can be skipped on a first reading.

The $\Delta^{(P)}$ operator is like a generalization of graph Laplacian. Indeed, when $P = 1$, the normal graph Laplacian $L$ is recovered. It turns out that the $\Delta^{(P)}$ operator defines a natural measure of smoothness for functions defined on the Hamming graph. In particular, for $P = 2$, one has

$$\epsilon_2^2(\phi) = \phi^T \Delta^{(2)} \phi = \sum_{f_{ij,kl}} \left[(\phi_i - \phi_j) - (\phi_k - \phi_l)\right]^2 \tag{10}$$

where $\mathrm{f}_{ij,kl}$ is the face (or "2-cube") formed by edge $\mathrm{e}_{ij}$ and edge $\mathrm{e}_{kl}$, and the corresponding epistatic coefficient is the (negative) conditional log odds ratio. Thus, the $\Delta^{(2)}$ operator measures the differences between parallel edges. It is intriguing to notice that these differences are actually the discrete analog of second-order mixed partial derivatives. Similarly, in the case of $P = 3$, one has

$$\epsilon_3^2(\phi) = \phi^T \Delta^{(3)} \phi = \sum_{\mathrm{c}_{ijkl,mnpq}} \Big\{ \big[ (\phi_i - \phi_j) - (\phi_k - \phi_l) \big] - \big[ (\phi_m - \phi_n) - (\phi_p - \phi_q) \big] \Big\}^2 \tag{11}$$

where $\mathrm{c}_{ijkl,mnpq}$ is the cube (or "3-cube") formed by face $\mathrm{f}_{ij,kl}$ and face $\mathrm{f}_{mn,pq}$. Again, the terms computed can be understood as the discrete analog of third-order mixed partial derivatives. In this work we focus on the operators $\Delta^{(2)}$ and $\Delta^{(3)}$ as they are closely related to the most widely used maximum entropy models and their interpretation is relatively straightforward.

#### Likelihood, prior, and posterior

Given a set of count data $\{N_i\}_{i=1}^G$ in the sequence space of interest, the likelihood of the data is given by

$$\mathcal{L} = \prod_{i=1}^{G} Q_i^{N_i} \tag{12}$$

where $Q_i$ is the probability of drawing the $i$th sequence. In order to be a valid probability distribution, the probabilities $\{Q_i\}_{i=1}^G$ must satisfy the constraints: $\sum_{i=1}^G Q_i = 1$ and $Q_i > 0$ for $i = 1, 2, \ldots, G$. We can parametrize the probabilities by a *field* $\phi$ as follows:

$$Q_i = \frac{e^{-\phi_i}}{\sum_{j=1}^G e^{-\phi_j}} \tag{13}$$

for $i = 1, 2, \ldots, G$. The advantage of doing such parametrization is that now both constraints on the probabilities are automatically fulfilled by construction and the field can take on any set of values.

In order to be able to control the smoothness of the probability distribution, we further impose the prior on $\phi$:

$$p(\phi|a) = \frac{1}{Z_a^0}\, e^{-S_a^0[\phi]}, \qquad S_a^0[\phi] = \frac{a}{2s} \sum_{i,j=1}^{G} \phi_i \Delta_{ij}^{(P)} \phi_j \tag{14}$$

where $a$ is a hyperparameter to be determined later, and $s$ is the number of $P$-cubes of the Hamming graph given by Eq. 5 and is a constant characteristic of the sequence space with a particular choice of the prior. The normalization constant of $p(\phi|a)$ is the *partition function* $Z_a^0$ given by

$$Z_a^0 = \int \mathcal{D}\phi\; e^{-S_a^0[\phi]} \tag{15}$$

namely, a functional integral of the *action* $S_a^0[\phi]$ over the field $\phi$ (one should note that since $\Delta^{(P)}$ has a nontrivial null space, this is in fact an improper prior and so Eq. 14 and Eq. 15 should be regarded as formal definitions). As described in the previous section, the $\Delta^{(P)}$ operator can measure the smoothness of functions defined on the graph, such as the field $\phi$ (or equivalently, the probability distribution $Q$). If $\phi$ is rough, $\phi^T \Delta^{(P)} \phi$ is large. For such $\phi$ to be preferable, the hyperparameter $a$ has to be small. Thus, the smoothness of the probability distribution is governed by the hyperparameter $a$.

Finally, multiplying the likelihood with the prior results in the posterior distribution of $\phi$:

$$p(\phi|\text{data}, a) = \frac{1}{Z_a}\, e^{-S_a[\phi]}, \qquad S_a[\phi] = \frac{a}{2s} \sum_{i,j=1}^{G} \phi_i \Delta_{ij}^{(P)} \phi_j + N \sum_{i=1}^{G} R_i \phi_i + N \sum_{i=1}^{G} e^{-\phi_i} \tag{16}$$

where $N = \sum_{i=1}^G N_i$ is the total count of data, $R_i = N_i/N$ for $i = 1, 2, \ldots, G$ is the observed frequency of the $i$th sequence, and the normalization constant $Z_a$ is given by the partition function

$$Z_a = \int \mathcal{D}\phi\; e^{-S_a[\phi]}. \tag{17}$$

Therefore, the posterior distribution of $\phi$ is a balance between two parts. One part tends to reproduce the data as much as possible, whereas the other part favors smoother $\phi$.

#### More on the prior

In this section we elaborate on the prior (Eq. 14), in particular, its meaning and properties. Later, we will provide a different, yet insightful, perspective of the prior.

As shown in Eqs. 2, 10, and 11, the $\Delta^{(P)}$ operator measures the smoothness of the field $\phi$ by looking at differences between its nodes, edges, and faces. Because of the parametrization in Eq. 13, differences between field values translate into logarithms of probability ratios (or odds), that is, $\phi_i - \phi_j = -\log(Q_i/Q_j)$, and differences between such logarithms in turn lead to logarithms of odds ratios, and so on. For example, when $P = 2$, the term associated with the face $\mathrm{f}_{ij,kl}$ is equal to

$$(\phi_i - \phi_j) - (\phi_k - \phi_l) = -\log \frac{(Q_i/Q_j)}{(Q_k/Q_l)} \tag{18}$$

which is the logarithm of an odds ratio. When $P = 3$, the term corresponding to the cube $\mathrm{c}_{ijkl,mnpq}$ becomes

$$\left[(\phi_i - \phi_j) - (\phi_k - \phi_l)\right] - \left[(\phi_m - \phi_n) - (\phi_p - \phi_q)\right] = -\log \frac{(Q_i/Q_j)/(Q_k/Q_l)}{(Q_m/Q_n)/(Q_p/Q_q)} \tag{19}$$

which is the logarithm of a ratio of odds ratios. Odds ratio, as the one in Eq. 18, is a common measure of the "association" (or correlation) between two quantities [3]. In our case, the four nodes forming the face $\mathrm{f}_{ij,kl}$ are sequences having two of the alleles at one site and having another two of the alleles at another site. For instance, they can be sequences with alleles (B,D), (A,D), (B,C), (A,C) at site 1 and site 2. To be clear, we can rewrite the odds ratio as (neglecting other sites)

$$-\log \frac{(Q_{\mathrm{BD}}/Q_{\mathrm{AD}})}{(Q_{\mathrm{BC}}/Q_{\mathrm{AC}})}. \tag{20}$$

Thus, the odds ratio here measures the particular association between the two sites, each with the specific change of allele (that is, A → B at site 1 and C → D at site 2). Similarly, the ratio of odds ratios in Eq. 19 measures the association between three different sites, where each site changes from one allele to another. In general, we have the following relation:

$$\phi^T \Delta^{(P)} \phi = \sum_{i=1}^{s} \left(\log \mathcal{A}_i^{(P)}\right)^2 \tag{21}$$

where $\mathcal{A}_i^{(P)} = \epsilon_{c^m_{b,A}}(\phi)$ is the association of $P$th order for the $P$-cube $i$ defined by $c^m_{b,A}$; for $P = 1,2,3$, it represents the conditional odds, odds ratio, and ratio of odds ratios, respectively. The summation is over all $P$-cubes (i.e., edges, faces, cubes, ...) for the computation of $\mathcal{A}^{(P)}$, and $s$ is the number of such $P$-cubes of the Hamming graph. Below we further show that the overall strength of the association is inversely proportional to the hyperparameter $a$.

The partition function $Z_a^0$ in Eq. 15 can be evaluated analytically. However, because $\Delta^{(P)}$ is a singular matrix, one should regularize the integral with a regularization parameter $\varepsilon$, namely,

$$Z_a^0 = \int \mathcal{D}\phi \; e^{-\frac{1}{2}\phi^T\left(\frac{a}{s}\Delta^{(P)} + \varepsilon I\right)\phi} \tag{22}$$

where $I$ is the identity matrix and $\varepsilon \to 0$. Carrying out the integral, one has

$$Z_a^0 = (2\pi)^{\frac{G}{2}} \left(\det\left[\frac{a}{s}\Delta^{(P)} + \varepsilon I\right]\right)^{-\frac{1}{2}} = (2\pi)^{\frac{G}{2}} \varepsilon^{-\frac{\kappa}{2}} a^{-\frac{G-\kappa}{2}} s^{\frac{G-\kappa}{2}} \left(\det_{\mathrm{row}}\left[\Delta^{(P)}\right]\right)^{-\frac{1}{2}} \tag{23}$$

where $\kappa$ is the kernel dimension of $\Delta^{(P)}$ and "row" represents the row space of $\Delta^{(P)}$. Differentiating Eq. 22 and Eq. 23 with respect to $a$ and then multiplying by $(-2)$, one obtains

$$(-2)\frac{dZ_a^0}{da} = \int \mathcal{D}\phi \left(\frac{1}{s}\,\phi^T \Delta^{(P)} \phi\right) e^{-\frac{1}{2}\phi^T\left(\frac{a}{s}\Delta^{(P)+\varepsilon I}\right)\phi} = \left(\frac{G-\kappa}{a}\right)(2\pi)^{\frac{G}{2}} \varepsilon^{-\frac{\kappa}{2}} a^{-\frac{G-\kappa}{2}} s^{\frac{G-\kappa}{2}} \left(\det_{\mathrm{row}}\left[\Delta^{(P)}\right]\right)^{-\frac{1}{2}}. \tag{24}$$

From these equations, we have the following simple relation:

$$\left\langle \frac{1}{s}\,\phi^T \Delta^{(P)} \phi \right\rangle_{p(\phi|a)} = \frac{G-\kappa}{a}. \tag{25}$$

In addition, from Eq. 21 the root-mean-square (rms) value of $\log \mathcal{A}^{(P)}$ can be expressed as follows:

$$\left(\log \mathcal{A}^{(P)}\right)_{\text{rms}} = \sqrt{\frac{1}{s}\sum_{i=1}^{s}\left(\log \mathcal{A}_i^{(P)}\right)^2} = \sqrt{\frac{1}{s}\,\phi^T \Delta^{(P)} \phi}\,. \tag{26}$$

Combining Eqs. 25 and 26, one gets the final result:

$$\left\langle \left(\log \mathcal{A}^{(P)}\right)^2_{\text{rms}} \right\rangle_{p(\phi|a)} = \frac{G-\kappa}{a}. \tag{27}$$

That is, the hyperparameter $a$ is inversely proportional to the expectation value of the squared rms $\log \mathcal{A}^{(P)}$ with respect to the prior distribution. Thus, by tuning $a$, one can control the overall association of the probability distribution $Q$.

#### Properties of the MAP solutions

Given a value of the hyperparameter $a$, the maximum a posteriori (MAP) estimate of the field, which corresponds to the mode of the posterior distribution $p(\phi|\text{data}, a)$ in Eq. 16, can be found by minimizing the action $S_a[\phi]$. Doing so, one can show that the MAP estimate, denoted $\phi^a$, satisfies the *equation of motion*:

$$\frac{a}{s}\sum_{j=1}^{G} \Delta_{ij}^{(P)} \phi_j^a + N R_i - N e^{-\phi_i^a} = 0 \tag{28}$$

for $i = 1, 2, \ldots, G$. With this equation of motion, one can derive some important properties of the MAP estimates.

Recall that the kernel of the operator $\Delta^{(P)}$ is given by $W^{(0)} \oplus W^{(1)} \oplus W^{(2)} \oplus \cdots \oplus W^{(P-1)}$, where $W^{(k)}$ is the subspace corresponding to the $k$th eigenvalue of the graph Laplacian $L$. Denote the basis of $W^{(k)}$ by $\{\psi_l^{(k)}\}_{l=1}^{m_k}$. Taking inner product of the equation of motion with the basis of $W^{(k)}$, one obtains

$$\sum_{i=1}^{G} \psi_{l,i}^{(k)} R_i = \sum_{i=1}^{G} \psi_{l,i}^{(k)} e^{-\phi_i^a} \tag{29}$$

for $l = 1, 2, \ldots, m_k$. For $k = 0$, the only basis vector is a constant vector of 1s. Thus, Eq. 29 leads to

$$\sum_{i=1}^{G} R_i = 1 = \sum_{i=1}^{G} e^{-\phi_i^a} \tag{30}$$

namely, the normalization factor in Eq. 13 for the MAP solution $\phi^a$ is 1 and, as a result, $Q_i^a = e^{-\phi_i^a}$. Therefore, Eq. 29 can be rewritten as

$$\sum_{i=1}^{G} \psi_{l,i}^{(k)} R_i = \sum_{i=1}^{G} \psi_{l,i}^{(k)} Q_i^a\,. \tag{31}$$

For $k \geq 1$, one can construct the basis in a specific way such that the inner product of the basis with a probability distribution will project out certain marginal frequencies of the probability distribution. For example, when $k = 1$, one can construct the basis by doing one-hot encoding to the array of all possible sequences, with each site being treated as a variable. Dropping the first allele at each site, this results in $m_1$ 1-site indicators. Using these indicators as the basis vectors can project out the marginal frequencies of having certain allele at certain site of the sequences. For $k = 2$, the basis can be constructed by forming product of the 1-site indicators from two different sites, repeated for all possible pairs of sites. This basis will consist of $m_2$ 2-site indicators and will be able to project out the marginal frequencies of having one allele at site 1 and another allele at site 2. This procedure can be continued to obtain higher-order marginal frequencies. Thus, given an operator $\Delta^{(P)}$, its kernel basis can be used to project out marginal frequencies involving up to $(P-1)$ sites of a probability distribution. Eq. 31 further indicates that these marginal frequencies of the MAP solution $Q^a$, for any $a$, is equal to the corresponding marginal frequencies of the observed frequency $R$. In other words, by choosing the order of the $\Delta^{(P)}$ operator, one can decide which marginal statistics of the data are to be preserved in the estimated probability distributions.

Moreover, in the limit of $a \to 0$, the equation of motion becomes

$$R_i = e^{-\phi_i^0} = Q_i^0 \, . \tag{32}$$

That is, the MAP solution of probability distribution in this limit is just the observed frequency. On the other hand, when $a \to \infty$, the MAP solution $\phi^\infty$ must satisfy

$$\sum_{j=1}^{G} \Delta_{ij}^{(P)} \phi_j^\infty = 0 \tag{33}$$

in order to make the equation of motion finite. Thus, the field $\phi^\infty$ must be a linear combination of the kernel basis of $\Delta^{(P)}$, and hence, is equal to the *maximum entropy* (MaxEnt) estimate. Depending on the order of the operator, and with the kernel basis constructed in the previous paragraph, the resulting probability distribution $Q^\infty$ can be additive MaxEnt (for $P = 2$), pairwise MaxEnt (for $P = 3$), and so forth. These two extreme cases of $a \to 0$ and $a \to \infty$ mark the two ends of the "MAP curve": on one end is the observed frequency with maximum association, on the other end is the MaxEnt estimate with minimum association. In between are probability distributions with various strengths of association, which increases monotonically as $a$ decreases. Given that every solution on the MAP curve, namely, $\{Q^a\}_{a=0}^{\infty}$, has the same projection on the kernel of $\Delta^{(P)}$, and that the projection on the kernel is just the MaxEnt solution, one can regard the MAP curve as a series of probability distributions starting as MaxEnt at $a \to \infty$, adding more and more structures as $a$ decreases, and eventually reaching the limit of the observed frequency at $a \to 0$.

#### Choosing the hyperparameter

The MAP curve consists of MAP estimates of the probability distribution $Q^a$, or equivalently the field $\phi^a$, for $a$ ranging from 0 to $\infty$. In practice, a finite set of $a$s, including 0 and $\infty$, is chosen to represent the whole MAP curve. This can be done by, for example, choosing the $a$s such that adjacent probability distributions are similar enough to each other. To find the optimal value of $a$, denoted $a^*$, one can either compute evidence ratio or do cross validation. Once $a^*$ is determined, the optimal probability distribution is given by $Q^* \equiv Q^{a^*}$. Similarly, $\phi^* \equiv \phi^{a^*}$.

The evidence ratio at $a$ is defined as the ratio of the Bayesian evidence at $a$ to the Bayesian evidence at $\infty$, that is, $E(a) = p(\text{data}|a)/p(\text{data}|\infty)$. It can be shown that the evidence ratio can be expressed in terms of four partition functions as the following:

$$E(a) = \frac{Z_a}{Z_a^0} \frac{Z_\infty^0}{Z_\infty} \tag{34}$$

where $Z_a^0$ and $Z_a$ respectively are the prior and posterior partition functions defined in Eq. 15 and Eq. 17, and $Z_\infty^0$ and $Z_\infty$ are the corresponding limits at $a \to \infty$. Except in some special cases, evaluating partition functions is a notoriously difficult task and one usually has to resort to some approximation, such as the Laplace approximation. Indeed, in such approximation, one can show that

$$\begin{aligned} \log E(a) = &\left\{ -S_a[\phi^a] + \frac{G-\kappa}{2} \log\left(\frac{a}{s}\right) - \frac{1}{2}\log\left(\det\left[\frac{a}{s}\Delta^{(P)} + Ne^{-\phi^a}\right]\right)\right\} \\ &- \left\{ -S_\infty[\phi^\infty] - \frac{1}{2}\log\left(\det_{\text{ker}}\left[Ne^{-\phi^\infty}\right]\right) - \frac{1}{2}\log\left(\det_{\text{row}}\left[\Delta^{(P)}\right]\right)\right\} \end{aligned} \tag{35}$$

where $\kappa$ is the kernel dimension of $\Delta^{(P)}$, and "ker" and "row" means the kernel and row space of $\Delta^{(P)}$, respectively. Corrections to this approximation can be estimated by using Feynman diagrams or sampling techniques [4]. Although this expression is analytic, it requires the determinant of several matrices. The first determinant in the second line of Eq. 35 is the determinant of the matrix mapped to the much smaller kernel of $\Delta^{(P)}$. Thus, its computation is not a problem. The second determinant in the same line is also easy to compute since the spectrum of $\Delta^{(P)}$ is analytically known. What causes problems is the determinant in the first line of Eq. 35. For the sequence spaces explored in this work, that is, human 5' splice sites and karyotypes of human cancer, the dimension of the space is from hundreds of thousands to several millions. Computing determinant of these large matrices is not feasible. One way to sidestep this difficulty is to find bounds on the determinant instead. Indeed, one basic property of determinant allows us to put a lower bound on the determinant in question, whereas an upper bound can be obtained by using the Hadamard's inequality [5]. Combining them altogether, the log-determinant term in the first line of Eq. 35 can be bounded as follows:

$$\sum_{i=1}^{G} \log\left(\frac{a}{s}\Delta_{ii}^{(P)} + Ne^{-\phi_i^a}\right) \geq \log\left(\det\left[\frac{a}{s}\Delta^{(P)} + Ne^{-\phi^a}\right]\right) \geq \sum_{i=1}^{G} \log\left(Ne^{-\phi_i^a}\right) \tag{36}$$

where $\Delta_{ii}^{(P)}$ is the $i$th diagonal element of $\Delta^{(P)}$ which is the same for all $i$. These inequalities in turn provide bounds on the evidence ratio. Empirically, we find that the lower bound in Eq. 36, which corresponds to the upper bound of the evidence ratio, is less informative. This is probably because the bound lacks information about the structure of the sequence space. In contrast, the upper bound in Eq. 36, which sets the lower bound of the evidence ratio, is found to be more informative and can sometimes result in values of $a^*$ that are consistent with the ones from cross validation (described below). However, whether this lower bound can be used as a surrogate for the evidence ratio itself requires further investigation.

Another approach to choose $a^*$ is by doing cross validation. This approach is more general in that it is not based on some approximation and is not restricted to small sequence spaces. In this work, we employ $k$-fold cross validation with $k = 5$. To implement it, the data is first split into $k$ subsets. Each subset is held out once while the remaining $(k-1)$ subsets are used to estimate MAP solutions $\{Q^a\}_{a=0}^{\infty}$. The log likelihood of the held-out subset is then computed with the resulting $Q^a$s. Finally, the averaged log likelihood over different held-out subsets is used to determine $a^*$. This procedure may seem very time-consuming as one has to repeat the same computations $k$ times, in addition to the original computations with the whole dataset. Nevertheless, this procedure can be readily parallelized and the $(k+1)$ MAP curves can be computed at the same time. In fact, one can also parallelize the computations at different $a$s of an MAP curve. Thus, the bottleneck of the whole computation is the slowest single MAP estimation, which is usually the one with the smallest (finite) $a$.

#### Posterior Sampling

To estimate the uncertainty in the resulting probability distribution or its derivatives, one can draw posterior samples from the joint posterior distribution $p(\phi, a|\text{data})$. Using Bayes' rule, we can rewrite $p(\phi, a|\text{data})$ as $p(\phi, a|\text{data}) = p(\phi|\text{data}, a)\, p(a|\text{data})$. The first term is the posterior distribution of $\phi$ given certain $a$, whereas the second term is the posterior distribution of $a$. We can further rewrite $p(a|\text{data})$ as $p(a|\text{data}) \propto p(\text{data}|a)\, p(a)$. Note that $p(\text{data}|a)$ is proportional to the evidence ratio $E(a)$ and $p(a)$ is usually assumed to be uniform over the $a$s. Therefore, the sampling procedure can be decomposed into two steps. First, a set of $a$s is sampled by using $E(a)$ [in the cases where $E(a)$ is not available, one can use the cross-validated likelihood as a substitute]. Then, for each $a$, we sample a number of $\phi$s from $p(\phi|\text{data}, a)$. Altogether, these $\phi$s at different $a$s form the posterior samples.

Whereas the first step of this sampling procedure is trivial, the second step is a challenging task. Again, this is because sequence spaces are usually very high-dimensional and sampling in such high-dimensional spaces will inevitably suffer from the curse of dimensionality. To mitigate the adversities, the method of Hamiltonian Monte Carlo (HMC) [6] exploits Hamiltonian dynamics to help explore the probability landscape. We outline HMC in the following. Imagine a particle moving in a $F$-dimensional space. The dynamics of the particle is governed by the *Hamiltonian*:

$$H(q, p) = U(q) + K(p) \tag{37}$$

where $q$ and $p$ respectively are the *position* and *momentum* of the particle, and $U$ is the *potential energy* shaping the landscape of the space while $K$ is the particle's *kinetic energy.* The trajectory of the particle is then determined by the Hamilton's equations:

$$\frac{dq_i}{dt} = \frac{\partial H}{\partial p_i}, \qquad \frac{dp_i}{dt} = -\frac{\partial H}{\partial q_i} \tag{38}$$

for $i = 1, 2, \ldots, F$. Here $t$ represents time. In the implementation of HMC, the potential energy $U$ is given by the negative logarithm of the probability distribution in question. In our case, $U = S_a[\phi]$, the posterior action, and so $q = \phi$. On the other hand, $p$ is an auxiliary vector and the kinetic energy $K$ is given by

$$K(p) = \frac{1}{2} \sum_{i,j=1}^{F} p_i M_{ij}^{-1} p_j \tag{39}$$

where $M$ is the *mass matrix*. In practice, $M$ is often set to a scalar multiple of the identity matrix. For each initial guess of $(q, p)$, we can evolve them with the Hamilton's equations using, for example, the leapfrog algorithm [6]. The resulting $(q^*, p^*)$ gives us a proposed state which is then accepted as the initial guess of next iteration with a probability of

$$\min\left(1, \frac{e^{-H(q^*,\, p^*)}}{e^{-H(q,\, p)}}\right). \tag{40}$$

If $(q^*, p^*)$ is not accepted, the original $(q, p)$ will serve as the initial guess again in next iteration. Hamiltonian dynamics can prevent the imaginary particle from sterile random walks in the space, and as a result, HMC has been found to be more efficient than other Monte Carlo techniques in high-dimensional problems.

In our practical applications of HMC to human 5' splice sites and karyotypes of human cancer, we found that having the stepsize properly scaled in each component is critical. In fact, it seemed infeasible to successfully implement HMC with a single common value of the stepsize $\varepsilon$. To estimate the scale in each component, we used the diagonal elements of the Hessian matrix:

$$(H_a)_{ij} \equiv \frac{\partial^2 S_a}{\partial \phi_i \partial \phi_j} = \frac{a}{s} \Delta_{ij}^{(P)} + N e^{-\phi_i^a} \delta_{ij}. \tag{41}$$

The stepsize in the $i$th component was then set to $\varepsilon_i = \varepsilon\, \sigma_i$, where $\sigma_i^{-2} = (H_a)_{ii}$. For simplicity, we only sampled at $a = a^*$. In each case, we ran 10 HMC chains. Each chain started with 1,000 warm-up draws. Thereafter, 100 samples were picked every 10 draws. In total, we generated 1,000 samples for each dataset. The stepsize parameter $\varepsilon$ was tuned to achieve an acceptance rate of about 0.65. We got $\varepsilon = 0.08$ for human 5' splice sites and $\varepsilon = 0.04$ for karyotypes of human cancer. On the other hand, the number of leapfrog steps had to be large enough to ensure mixing among different HMC chains. To assess the degree of mixing, we employed the statistic $\hat{R}$ [7], which is an estimate of the ratio of between-chain variance to within-chain variance. A value of $\hat{R}$ close to 1 indicates that the HMC chains are well mixed. We used 25 leapfrog steps for human 5' splice sites and 75 leapfrog steps for karyotypes of human cancer. This resulted in $\hat{R} = 0.991 \sim 1.023$ and $\hat{R} = 0.992 \sim 1.147$ for the two datasets, respectively.

### A Different Perspective on the Prior

The prior given in Eq. 14 has been interpreted as a constraint on the field $\phi$ which enforces the corresponding probability distribution $Q$ to have an expected strength of local associations dictated by the hyperparameter $a$ through Eq. 27. In this interpretation, one has to compute the measure of the association for each $P$-cube of the Hamming graph with a given $\phi$. For example, for $P = 2$, the $P$-cube is the face formed by four sequences and the associated measure is the odds ratio. The root-mean-square value of the measure, together with the hyperparameter $a$, then determines the probability of such $\phi$. The prior can, in fact, also be interpreted the other way around. Specifically, the association measure of each $P$-cube can be regarded to follow a zero mean normal distribution with variance governed by the hyperparameter $a$, and with each set of the association measures of all the $P$-cubes sampled from this distribution one can attempt to "reconstruct" a field $\phi$ that would produce the sampled local associations. Such $\phi$ will then have an expected overall association determined by $a$. This interpretation is particularly helpful in comparison with the associated MaxEnt estimate which sets the association for each $P$-cube to zero, since SeqDEFT prior can be viewed as replacing this deterministic zero value with draws from a zero mean Gaussian.

More precisely, given a field $\phi \in \mathbb{R}^G$, one can extract the $P$th-order epistatic coefficient $\epsilon$ for all $P$-cubes of the Hamming graph using a $s \times G$ matrix $C$ as follows:

$$\epsilon = C\phi\,. \tag{42}$$

From Eq. 8 it is clear that

$$C^T C = \Delta^{(P)}. \tag{43}$$

We call a vector $\epsilon \in \mathbb{R}^s$ compatible if there exists a field $\phi \in \mathbb{R}^G$ such that $C\phi = \epsilon$. Recall that the kernel of $\Delta^{(P)}$ is given by $W^{(0)} \oplus W^{(1)} \oplus W^{(2)} \oplus \cdots \oplus W^{(P-1)}$, where $W^{(k)}$ is the subspace corresponding to the $k$th eigenvalue of the graph Laplacian $L$. On the other hand, for $k \geq P$, one has $\Delta^{(P)} W^{(k)} = W^{(k)}$. Thus, the range of $\Delta^{(P)}$ is equal to $W^{(P)} \oplus W^{(P+1)} \oplus \cdots \oplus W^{(\ell)}$, and the space of $\mathbb{R}^G$ can be decomposed as

$$\mathbb{R}^G = \ker\big(\Delta^{(P)}\big) \oplus \operatorname{range}\big(\Delta^{(P)}\big). \tag{44}$$

Since $W^{(k)}$ corresponds to the space of all $k$th-order interactions, for any $\phi \in \mathbb{R}^G$, we can decompose it into two parts, namely, $\phi = \phi_{<P} + \phi_{\geq P}$, where $\phi_{<P} \in \ker\big(\Delta^{(P)}\big)$ is the component having interactions of order lower than $P$ and $\phi_{\geq P} \in \operatorname{range}\big(\Delta^{(P)}\big)$ is the component having interactions of order equal to and higher than $P$. Define the projection operator $\Pi : \phi \mapsto \phi_{\geq P}$. We can project the prior distribution $p(\phi|a)$ in Eq. 14 with $\Pi$ to obtain a marginal distribution over the space of all epistatic components:

$$p_\Pi(\phi|a) \equiv \Pi\big(p(\phi|a)\big) = \frac{1}{Z}\, e^{-\frac{a}{2s}\phi^T \Delta^{(P)} \phi} \tag{45}$$

with $\phi \in \operatorname{range}\big(\Delta^{(P)}\big)$. The normalization constant $Z$ is equal to

$$Z = \sqrt{\det\left[\frac{2\pi s}{a} \Delta^{(P)^+}\right]} \tag{46}$$

where $\Delta^{(P)^+}$ is the Moore-Penrose pseudoinverse of $\Delta^{(P)}$ and its determinant is given by the product of the nonzero eigenvalues. The distribution $p_\Pi(\phi|a)$ has the same form as the prior distribution $p(\phi|a)$, however, $p_\Pi(\phi|a)$ is defined over the subspace $\operatorname{range}\big(\Delta^{(P)}\big)$, whereas $p(\phi|a)$ is defined over the space $\mathbb{R}^G$. Now we state the main result of this section.

**Proposition 2.** *If we sample the entries of $\epsilon$ independently from a normal distribution with mean 0 and variance $\frac{s}{a}$ while conditioning on $\epsilon$ being compatible, the corresponding random field $\phi$ follows the same distribution as Eq. 45.*

The proof of this proposition is provided in Appendix 2.

### Computing Pairwise Association

Once an estimate of probability distribution in a sequence space is obtained, such as $Q^*$, there is a wealth of information regarding the intrinsic nature of the sequence space that can be extracted. For example, the associations between different sites of the sequences is very informative as they are manifestations of the underlying biological mechanism. Knowing the manner in which the sites associate can therefore help one figure out the related biology. There are many kinds of associations, each involves a different number of sites. In this work we focus on "pairwise association", that is, association between two sites.

The strength of the association between two quantities $X$ and $Y$ can be quantified by the odds ratio [3]. Assuming that $X$ and $Y$ can both be 0 or 1, the odds ratio is given by

$$\frac{(p_{11}/p_{01})}{(p_{10}/p_{00})} \tag{47}$$

where $p_{xy}$ represents the probability of $X = x$ and $Y = y$. The odds ratio is the ratio of the odds of $X$ conditional on $Y = 1$ to the odds of $X$ conditional on $Y = 0$. If the odds ratio is 1, namely, the odds of $X$ is the same no matter what $Y$ is, it means that $X$ and $Y$ are not associated. If the odds ratio is greater than 1, it means that $X$ has a greater odds when $Y = 1$ than when $Y = 0$. In this case, $X$ and $Y$ are called positively associated, and the larger the odds ratio, the stronger the association. Similarly, if the odds ratio is less than 1, $X$ and $Y$ are said to be negatively associated. We can use some numbers to illustrate this concept. If $X$ and $Y$ are not associated, it means we will observe the same amount (say, 50 and 50) of 1s and 0s for $X$ whether $Y = 1$ or $Y = 0$. If $X$ and $Y$ are positively associated, we will observe a lot of 1s and a few of 0s (say, 90 and 10) for $X$ when $Y = 1$, but we will observe a few of 1s and a lot of 0s (say, 10 and 90) when $Y = 0$. For the case where $X$ and $Y$ are negatively associated, the numbers above are switched. Thus, odds ratio is a measure of association, or correlation, and is symmetric with respect to the two quantities. However, it is important to notice that association does not imply causality.

In our case, in order to compute association between two sites of the sequences, one has to specify the "mutation" at each site. For example, the first site changes from allele A to allele C and the second site changes from allele G to allele T. The corresponding odds ratio is defined as (neglecting other sites)

$$\frac{(Q^*_{\mathrm{CT}}/Q^*_{\mathrm{AT}})}{(Q^*_{\mathrm{CG}}/Q^*_{\mathrm{AG}})}. \tag{48}$$

So the association here is particular about the specific mutations at the two sites. For $\alpha$ alleles, there are $\binom{\alpha}{2}^2$ such combinations of mutations at two sites. For each combination of mutations at two sites, there are $\alpha^{\ell-2}$ arrangements of the remaining sites, or "backgrounds". Thus, for each specification of mutations at two sites, one actually obtains a distribution of the odds ratio. Depending on the underlying biology, the distribution may have certain structure, such as being normal or skewed, or having one or multiple modes. This information can also allow one to find the backgrounds where a particular association is strongest.

### Visualizing Probability Landscape

Sequence space is generally a very high-dimensional space. To "see" functions in such space is therefore very challenging. Nevertheless, there exist various techniques of dimensionality reduction that allow one to visualize such high-dimensional objects by projecting them into a lower dimensional space. In this work we adapt the method developed in [8] to visualize the resulting probability distribution $Q^*$. These visualizations can reveal structures of the probability landscape. From the geometry of the sequences, useful information regarding the underlying biology can also be obtained.

To make visualization of the probability distribution $Q^*$, we first must build a transition (or rate) matrix $T$ for a reversible Markov chain with stationary distribution given by $Q^*$ that makes local transitions in the sequence space. The transition matrix we employ here is given by

$$T_{ij} = \begin{cases} \frac{\log Q^*_j - \log Q^*_i}{1 - e^{-(\log Q^*_j - \log Q^*_i)}} & \text{if } i \neq j \text{ and } d(\sigma_i, \sigma_j) = 1 \\ 0 & \text{if } i \neq j \text{ and } d(\sigma_i, \sigma_j) \neq 1 \\ -\sum_{k=1, k\neq i}^{G} T_{ik} & \text{if } i = j \end{cases} \tag{49}$$

for $i, j = 1, 2, \ldots, G$, where $d(\sigma_i, \sigma_j)$ is the Hamming distance between sequence $\sigma_i$ and sequence $\sigma_j$. There are many other choices of the transition matrix that have the stationary distribution $Q^*$. Empirically, it is found that different choices of transition matrix lead to similar results so long as the resulting Markov chain is reversible and the $T_{ij}$s are roughly order 1 for adjacent $(i, j)$. (Intuitively, this makes the major features of the Markov chain correspond to the global rather than local geometry of the probability distribution, as opposed to rates defined as e.g. $T_{ij} = Q^*_{ij}$.) Define $D_x$ to be the diagonal matrix with the vector $x$ down its main diagonal. The transition matrix can be symmetrized as follows:

$$\widetilde{T} = D_{Q^*}^{\frac{1}{2}}\, T\, D_{Q^*}^{-\frac{1}{2}}. \tag{50}$$

Choose $c$ from $\widetilde{T}$ such that $c = \max_i \sum_{j=1}^{G} |\widetilde{T}_{ij}|$, and then let

$$A = I + \frac{1}{c}\widetilde{T} \tag{51}$$

where $I$ is the identity matrix. Compute the dominant eigenvalues $1 = \widetilde{\lambda}_0 \geq \widetilde{\lambda}_1 \geq \ldots$ of the symmetric matrix $A$ with associated orthonormal eigenvectors $\widetilde{u}_0, \widetilde{u}_1, \ldots$. Then the $k$th largest eigenvalue of $T$ is $\lambda_k = c\,(\widetilde{\lambda}_k - 1)$

and the associated $D_{Q^*}$-orthonormal right eigenvector of $T$ is $u_k = D_{Q^*}^{-\frac{1}{2}}\,\widetilde{u}_k$. A $k$-dimensional visualization can be done by plotting the $i$th sequence at coordinates:

$$\left(\frac{1}{\sqrt{-\lambda_1}}\,u_{1,i}, \frac{1}{\sqrt{-\lambda_2}}\,u_{2,i}, \ldots, \frac{1}{\sqrt{-\lambda_k}}\,u_{k,i}\right) \tag{52}$$

for $i = 1, 2, \ldots, G$. The factor $1/(-\lambda_k)$ gives the fraction of variance explained in the $k$th direction. Also, when $k = G$, the "commute time" between sequence $\sigma_i$ and sequence $\sigma_j$, that is, the time traveling from $\sigma_i$ to $\sigma_j$ plus the time traveling from $\sigma_j$ to $\sigma_i$, is proportional to the squared Euclidean distance between $\sigma_i$ and $\sigma_j$ in the high-dimensional representation. For $k < G$, the result is equivalent to doing weighted principal component analysis on this high-dimensional representation when sequences are weighted by their probabilities under $Q^*$.

### Appendix 1: Proof of Proposition 1

From the definition in Eq. 6 and Eq. 7, we note that $\epsilon_P^2(\phi)$ consists of the sum of squared polynomials, so that $\epsilon_P^2(\phi) \geq 0$, and because the objects being squared are homogenous polynomials of degree 1, $\epsilon_P^2(\phi)$ is a homogeneous polynomial of degree 2. Thus we know that we can express $\epsilon_P^2(\phi)$ as a positive semi-definite quadratic form. That is, we can define a matrix $\Delta^{(P)}$ such that

$$\epsilon_P^2(\phi) = \phi^T \Delta^{(P)} \phi\,. \tag{53}$$

Our goal here is to show that $\Delta^{(P)}$ is actually given by $\frac{1}{P!}L^{(P)}$.

We begin by deriving the entries of $\Delta^{(P)}$. Expanding $\epsilon_P^2(\phi)$ using Eq. 7 and Eq. 53, we have

$$\epsilon_P^2(\phi) = \sum_{\sigma_i,\sigma_j} (-1)^{h(\sigma_i')+h(\sigma_j')} n(\sigma_i,\sigma_j)\phi(\sigma_i)\phi(\sigma_j) = \sum_{\sigma_i,\sigma_j} \Delta^{(P)}(\sigma_i,\sigma_j)\phi(\sigma_i)\phi(\sigma_j) \tag{54}$$

where $n(\sigma_i,\sigma_j)$ is the number of $P$-cubes in which $\sigma_i$ and $\sigma_j$ co-occur. For a pair of sequences $\sigma_i$ and $\sigma_j$ that are separated by a Hamming distance $d$, a $P$-cube containing both sequences can be constructed by first choosing $(P-d)$ sites among the $(\ell-d)$ background sites, and then for each of the $(P-d)$ sites introducing an alternative allele to form a complete $P$-cube. Therefore, the number of $P$-cubes containing both sequences is equal to $n(\sigma_i,\sigma_j) = \binom{\ell-P}{P-d}(\alpha-1)^{P-d}$. Now within such a hypercube, we can arbitrarily set $\sigma_i'$ to be the all-0 sequence, and thereby, $(-1)^{h(\sigma_i')+h(\sigma_j')} = (-1)^d$. Putting these together, we find

$$\Delta^{(P)}(\sigma_i,\sigma_j) = \begin{cases} (-1)^d\binom{\ell-d}{P-d}(\alpha-1)^{P-d} & \text{if } d(\sigma_i,\sigma_j) \leq P \\ 0 & \text{otherwise} \end{cases}. \tag{55}$$

Note that $\Delta^{(P)}(\sigma_i,\sigma_j) = (-1)^P$ if the Hamming distance between the two sequences is $P$, since in this case there is exactly one $P$-cube that contains both sequences. Importantly, given $\ell$, $\alpha$, and $P$, $\Delta^{(P)}(\sigma_i,\sigma_j)$ is a function of the Hamming distance $d$ only and has at most $(\ell+1)$ distinct values—a property shared by $L$ and its powers. In fact, $\Delta^{(P)}$ can be written as a polynomial in $L$ as follows [9]:

$$\Delta^{(P)} = \sum_{k=0}^{P} b_k L^k. \tag{56}$$

To see this, recall that $L = D - A$, where $D$ is the degree matrix which is equal to a multiple of the identity matrix due to the regularity of the Hamming graph, and $A$ is the adjacency matrix which has entry 1 if the corresponding sequences are adjacent and 0 otherwise. We then find

$$L^k = (-1)^k A^k + \sum_{i=1}^{k-1}\binom{k}{i} D^{k-i}(-A)^i. \tag{57}$$

It is well known that $A^n(\sigma_i,\sigma_j)$ is equal to the number of paths of length $n$ that start at $\sigma_i$ and end at $\sigma_j$. Thus, $A^n(\sigma_i,\sigma_j) = n!$ if $d(\sigma_i,\sigma_j) = n$, and $A^n(\sigma_i,\sigma_j) = 0$ if $d(\sigma_i,\sigma_j) > n$. This implies that

$$L^k(\sigma_i,\sigma_j) = (-1)^k k! \tag{58}$$

if $d(\sigma_i, \sigma_j) = k$, and $L^k(\sigma_i, \sigma_j) = 0$ if $d(\sigma_i, \sigma_j) > k$. Since $L^k$ assumes at most $(\ell + 1)$ distinct values as a function of Hamming distance, we can form a $(\ell+1) \times (\ell+1)$ matrix whose $k$th column contains entries of $L^{k-1}$ arranged by Hamming distance. The resulting matrix is upper triangular with nonzero entries along the main diagonal. Thus, $L^0, L^1, L^2, \ldots, L^\ell$ are linearly independent when viewed as vectors, and any matrix whose entries are a function of Hamming distance can be represented as a polynomial in $L$. Moreover, according to Eq. 55, $\Delta^{(P)}$ must be a polynomial in $L$ of degree $P$, which justifies Eq. 56.

Next, note that the kernel of $L^{(P)}$ is given by

$$\ker\big(L^{(P)}\big) = W^{(0)} \oplus W^{(1)} \oplus W^{(2)} \oplus \cdots \oplus W^{(P-1)} \equiv Y \tag{59}$$

where $W^{(k)}$ is the subspace corresponding to the $k$th eigenvalue of the graph Laplacian $L$. We now show that the matrix $\Delta^{(P)}$ also has $Y$ as its kernel. First, we construct a set of vectors that span $Y$ in the following manner. Let $r \subset \{1, 2, \ldots, \ell\}$ be a subset of size $k < P$ and $\mu$ a sub-sequence of length $k$. For a sequence $\sigma$, we can define the indicator function

$$\psi^r_\mu(\sigma) = \prod_{i=1}^{k} \delta_{\,\sigma(r(i)),\, \mu(i)} \tag{60}$$

where $\delta$ is the Kronecker delta. This indicator function gives 1 if the sequence $\sigma$ has the motif specified by $\mu$ at the sites $r$, and 0 otherwise. The functions

$$\Big\{\psi^r_\mu : r \subset \{1, 2, \ldots, \ell\},\ |r| < P\ ;\ \mu \in \{1, 2, \ldots, \alpha\}^k,\ k < P\Big\} \tag{61}$$

then form a spanning set for the space $Y$. For a function $\psi^r_\mu$ and a $P$-cube $c^m_{b,A}$, we can choose a site $i \in m \setminus r \neq \varnothing$ and let $\sigma^{(i)}$ be the sequence that differ from $\sigma$ at site $i$. Then, one has $\psi^r_\mu(\sigma) = \psi^r_\mu(\sigma^{(i)})$. Since the $P$-cube $c^m_{b,A}$ can be divided into pairs of sequences like these, and $|h(\sigma^{(i)\prime}) - h(\sigma')| = 1$, Eq. 6 reduces to

$$\epsilon_{c^m_{b,A}}(\psi^r_\mu) = \frac{1}{2} \sum_{\sigma \in c^m_{b,A}} \Big((-1)^{h(\sigma')} + (-1)^{h(\sigma')+1}\Big)\psi^r_\mu(\sigma) = 0\,. \tag{62}$$

This procedure can be repeated for every $P$-cube, and thus, we have $\psi^r_\mu \in \ker\big(\Delta^{(P)}\big)$. On the other hand, we can also define indicator functions $\psi^t_\nu$ as in Eq. 60 but with the size of the subset $t$ equal to or greater than $P$. In this case, we can first find one sequence with the motif $\nu$ on sites $t$, and then construct a $P$-cube $c^m_{b,A}$ such that $m \subset t$. It is clear that there is only one sequence with the correct motif in $c^m_{b,A}$. As a result, we find $\epsilon^2_{c^m_{b,A}}(\psi^t_\nu) = 1$. Consequently,

$$\psi^{t\,T}_\nu \Delta^{(P)} \psi^t_\nu > 0 \tag{63}$$

which means that $\psi^t_\nu \notin \ker\big(\Delta^{(P)}\big)$. Thus, we have $\ker\big(\Delta^{(P)}\big) = Y$.

Finally, let $X$ denote the set of matrices

$$X = \left\{B = \sum_{k=0}^{P} b_k L^k : b_P \neq 0 \text{ and } \ker(B) = Y\right\}. \tag{64}$$

Our results so far have established that both $\Delta^{(P)}$ and $L^{(P)}$ belong to $X$. We now show that $X$ is a one-dimensional linear subspace, so these two matrices only differ by a multiplicative constant. For every $B \in X$, its eigenvalues must satisfy:

$$\lambda_i(B) = \sum_{k=0}^{P} b_k (i\alpha)^k = 0 \tag{65}$$

for $i = 0, 1, 2, \ldots, P-1$. Or equivalently, the coefficients $b_k$ must satisfy the linear equation:

$$\begin{bmatrix} 1 & 0 & 0 & \cdots & 0 \\ 1 & \alpha & \alpha^2 & \cdots & \alpha^P \\ 1 & 2\alpha & (2\alpha)^2 & \cdots & (2\alpha)^P \\ \vdots & \vdots & \vdots & & \vdots \\ 1 & (P-1)\alpha & ((P-1)\alpha)^2 & \cdots & ((P-1)\alpha)^P \end{bmatrix} \begin{bmatrix} b_0 \\ b_1 \\ b_2 \\ \vdots \\ b_P \end{bmatrix} = \begin{bmatrix} 0 \\ 0 \\ 0 \\ \vdots \\ 0 \end{bmatrix}. \tag{66}$$

This implies that $b_0 = 0$. Therefore, this equation reduces to

$$\begin{bmatrix} \alpha & \alpha^2 & \cdots & \alpha^{P-1} \\ 2\alpha & (2\alpha)^2 & \cdots & (2\alpha)^{P-1} \\ \vdots & \vdots & & \vdots \\ (P-1)\alpha & ((P-1)\alpha)^2 & \cdots & ((P-1)\alpha)^{P-1} \end{bmatrix} \begin{bmatrix} b_1 \\ b_2 \\ \vdots \\ b_{P-1} \end{bmatrix} = -b_P \begin{bmatrix} \alpha^P \\ (2\alpha)^P \\ \vdots \\ ((P-1)\alpha)^P \end{bmatrix}. \tag{67}$$

Since $b_P \neq 0$ by the definition of $X$, we can re-express the equation as:

$$\begin{bmatrix} \alpha & \alpha^2 & \cdots & \alpha^{P-1} \\ 2\alpha & (2\alpha)^2 & \cdots & (2\alpha)^{P-1} \\ \vdots & \vdots & & \vdots \\ (P-1)\alpha & ((P-1)\alpha)^2 & \cdots & ((P-1)\alpha)^{P-1} \end{bmatrix} \begin{bmatrix} \frac{b_1}{b_P} \\ \frac{b_2}{b_P} \\ \vdots \\ \frac{b_{P-1}}{b_P} \end{bmatrix} \equiv M\widetilde{b} = -\begin{bmatrix} \alpha^P \\ (2\alpha)^P \\ \vdots \\ ((P-1)\alpha)^P \end{bmatrix}. \tag{68}$$

Dividing each row of the matrix $M$ by its leading entry, we obtain a Vandermonde matrix:

$$V = \begin{bmatrix} 1 & \alpha & \cdots & \alpha^{P-2} \\ 1 & 2\alpha & \cdots & (2\alpha)^{P-2} \\ \vdots & \vdots & & \vdots \\ 1 & (P-1)\alpha & \cdots & ((P-1)\alpha)^{P-2} \end{bmatrix}. \tag{69}$$

By the properties of Vandermonde matrices, $\det(V) = \prod_{1\le i<j\le P-1}(j\alpha - i\alpha) \neq 0$. Since the matrix $M$ has rows that are multiples of the rows of $V$, we have $\det(M) = (P-1)!\,\alpha^{P-1}\det(V) \neq 0$. Therefore, Eq. 68 has a unique solution and the matrix $B$ can be determined up to a multiplicative constant, that is,

$$X = \left\{ c\left(\sum_{k=1}^{P-1}\widetilde{b}_k L^k + L^P\right) : c \in \mathbb{R} \right\} \tag{70}$$

where $c$ is a constant and $\widetilde{b}$ is the solution of Eq. 68. In other words, $X$ is a one-dimensional subspace, and thereby,

$$\Delta^{(P)} = c^* \, L^{(P)} \tag{71}$$

for some constant $c^*$. Since the leading coefficient for $\Delta^{(P)}$ is equal to $(-1)^P$ according to Eq. 55, while that of $L^{(P)}$ is $(-1)^P P!$ by Eq. 58, we find $c^* = \frac{1}{P!}$, which completes the proof.

### Appendix 2: Proof of Proposition 2

From Eq. 42, the set of all compatible $\epsilon$s form a subspace of $\mathbb{R}^s$ which is equal to range($C$). It can be shown that range($C$) = range($CC^+$)[1], where $C^+$ is the Moore-Penrose pseudoinverse of $C$. The singular value decomposition (SVD) of $C$ and $C^+$ can be expressed as $C = U\Sigma V^T$ and $C^+ = V\Sigma^+U^T$, respectively. This gives us

$$CC^+ = U\Sigma V^T V\Sigma^+ U^T = U\Sigma\Sigma^+U^T. \tag{72}$$

Since $U$ is orthonormal and $\Sigma\Sigma^+$ is a diagonal matrix, $\Sigma\Sigma^+$ contains the eigenvalues of $CC^+$, including $m = \text{rank}(C)$ 1s and $(s-m)$ 0s. We can write $U$ as $U = [Q, R]$, where the columns of $Q$ correspond to the nonzero eigenvalues and form an orthonormal basis for range($CC^+$) while the columns of $R$ form a basis for the orthogonal complement. One can verify that $QQ^T = CC^+$.

Next, let $\epsilon \in \mathbb{R}^s$ be a random variable following the distribution $q = \mathcal{N}(0, \frac{s}{a}I_s)$. Since we wish to sample only the compatible $\epsilon$ but the distribution $q$ we just defined has support over the whole space $\mathbb{R}^s$, we have to work with the conditional distribution given the event that $\epsilon$ is compatible. Briefly speaking, this is equivalent to projecting the distribution $q$ to the subspace range($CC^+$), which will give us a normal distribution with

[1] We prove range($C$) = range($CC^+$) by showing that (i) range($C$) $\subset$ range($CC^+$), and (ii) range($CC^+$) $\subset$ range($C$).

(i) For every $w \in$ range($C$), there exists a vector $\phi \in \mathbb{R}^G$ such that $w = C\phi$. By the property of the pseudoinverse, we have $w = C\phi = (CC^+)C\phi \in$ range($CC^+$).

(ii) For every $v \in$ range($CC^+$), there exists a vector $\eta \in \mathbb{R}^s$ such that $v = (CC^+)\eta = C(C^+\eta) \in$ range($C$).

zero mean and covariance matrix $CC^+CC^+ = CC^+$. Specifically, we can transform the random variable $\epsilon$ to a new coordinate using the orthonormal matrix $U$, that is,

$$\epsilon' = U^T \epsilon = \begin{bmatrix} Q^T \\ R^T \end{bmatrix} \epsilon = \begin{bmatrix} \epsilon_1' \\ \epsilon_2' \end{bmatrix}. \tag{73}$$

One can show that the transformed variable $\epsilon'$ follows the original distribution $q$ as follows:

$$q'(\epsilon') = q(U\epsilon') \cdot 1 \propto e^{-(U\epsilon')^T U\epsilon'} = e^{-(\epsilon')^T \epsilon'}. \tag{74}$$

It is easy to verify that $e \in \text{range}(CC^+)$ if and only if $R^T e = e_2' = 0$. Therefore,

$$q(\epsilon|\epsilon \in \text{range}(CC^+)) = q(\epsilon_1'|\epsilon_2' = 0) = \frac{1}{Z}\, e^{-\frac{a}{2s}\epsilon_1'^T \epsilon_1'} = \frac{1}{Z}\, e^{-\frac{a}{2s}\epsilon^T QQ^T \epsilon} = \frac{1}{Z}\, e^{-\frac{a}{2s}\epsilon^T CC^+ \epsilon}. \tag{75}$$

That is, the conditional distribution $q(\epsilon|\epsilon \in \text{range}(CC^+))$ is a normal distribution with covariance matrix $\frac{s}{a}CC^+$.

Having defined the distribution over the space of all compatible $\epsilon$s, we now proceed to define the distribution over the space of all possible $\phi$s constructed from such $\epsilon$s. Specifically, given a compatible $\epsilon$, we have $\phi = C^+\epsilon$. Using the property of the pseudoinverse, one finds that $C\phi = CC^+\epsilon = CC^+(CC^+\eta) = CC^+\eta = \epsilon$, with some vector $\eta \in \mathbb{R}^s$. In other words, we can naturally find a vector $\phi$ for any compatible $\epsilon$ such that the epistatic coefficients of $\phi$ is equal to $\epsilon$. It is easy to verify that the space of all $\phi$s constructed using this method is $\text{range}(C^+CC^+) = \text{range}(C^+) = \text{range}(C^+C)$, where the last equality is achieved following the same argument as 1. Also, we have $C^+C = V\Sigma^+U^TU\Sigma V^T = V\Sigma^+\Sigma V^T$ and $\Delta^{(P)} = C^TC = V\Sigma^TU^TU\Sigma V^T = V\Sigma^T\Sigma V^T$. Since both $\Sigma^+\Sigma$ and $\Sigma^T\Sigma$ are diagonal, $C^+C$ and $\Delta^{(P)}$ can be simultaneously diagonalized by $V$. Furthermore, since $\Sigma^+\Sigma$ and $\Sigma^T\Sigma$ have nonzero entries in the same positions along the main diagonal, we find that $\text{range}\big(\Delta^{(P)}\big) = \text{range}(C^+C) = \text{range}(C^+)$.

For a random variable $\epsilon$ that is normally distributed as in Eq. 75, the linearly transformed variable $\phi = C^+\epsilon$ is also normally distributed. According to the previous paragraph, the distribution of $\phi$ has support only in $\text{range}(C^+) = \text{range}\big(\Delta^{(P)}\big)$ and can be written as

$$r(\phi) = \frac{1}{Z'}\, e^{-\frac{1}{2}\phi^T K^+ \phi} \tag{76}$$

for $\phi \in \text{range}\big(\Delta^{(P)}\big)$, with the covariance matrix $K$ given by

$$K = \mathbb{E}\left[\phi\phi^T\right] = \mathbb{E}\left[C^+\epsilon(C^+\epsilon)^T\right] = C^+\,\mathbb{E}\left[\epsilon\epsilon^T\right](C^+)^T = \frac{s}{a}\,C^+CC^+(C^+)^T = \frac{s}{a}\,C^+(C^+)^T. \tag{77}$$

Here we have used the pseudoinverse of $K$ since $K$ can be singular. Finally, using the SVD of $C^+$, we find

$$\begin{aligned}\frac{s}{a}\,K^+ &= (C^+(C^+)^T)^+ = (V\Sigma^+(\Sigma^+)^TV^T)^+ = V(\Sigma^+(\Sigma^+)^T)^+V^T = V((\Sigma^+)^T)^+(\Sigma^+)^+V^T \\ &= V\Sigma^T\Sigma V^T = V\Sigma^TU^TU\Sigma V^T = C^TC = \Delta^{(P)}.\end{aligned} \tag{78}$$

Plugging this to Eq. 76, we find that the distribution $r(\phi)$ is identical to $p_\Pi(\phi|a)$ in Eq. 45.

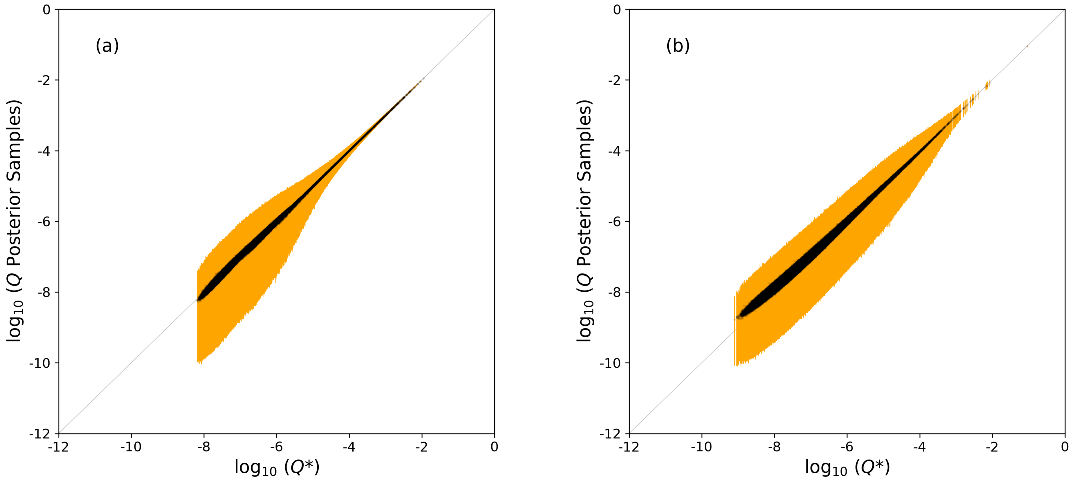


**Figure 1:** Mean (black dot) and 95% credible interval (orange line) of the probability distribution $Q$ (estimated from of 1,000 posterior samples) versus the optimal estimate $Q^*$ using $a^*$ chosen to maximimize the cross-validated log likelihood for (a) human 5' splice sites and (b) karyotypes of human cancer.

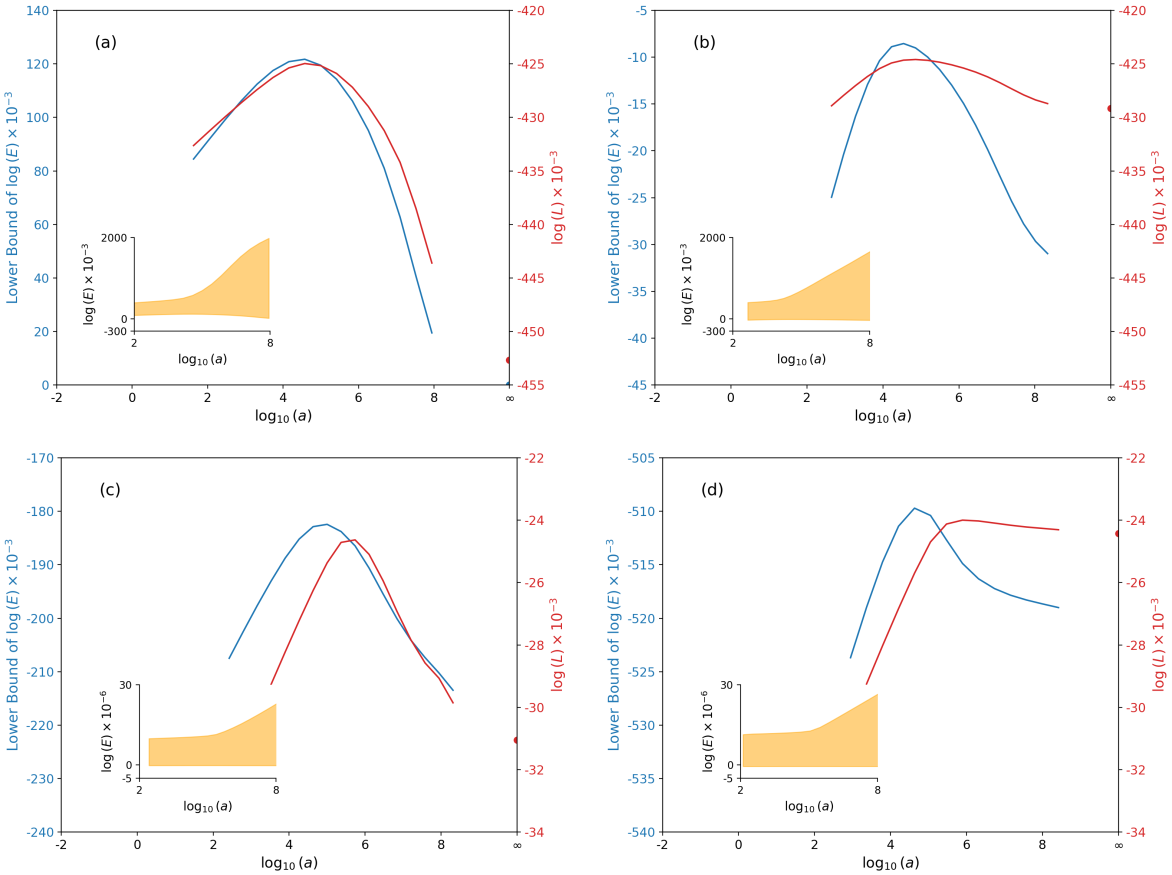


**Figure 2:** Lower bound of evidence ratio ($E$) and likelihood from cross validation ($L$) versus hyperparameter $a$ for human 5' splice sites [(a) for $P = 2$ and (b) for $P = 3$] and karyotypes of human cancer [(c) for $P = 2$ and (d) for $P = 3$] computed by SeqDEFT. Shown in the insets is the full range of $E$ versus $a$ from the bounds given in Eq. 36.

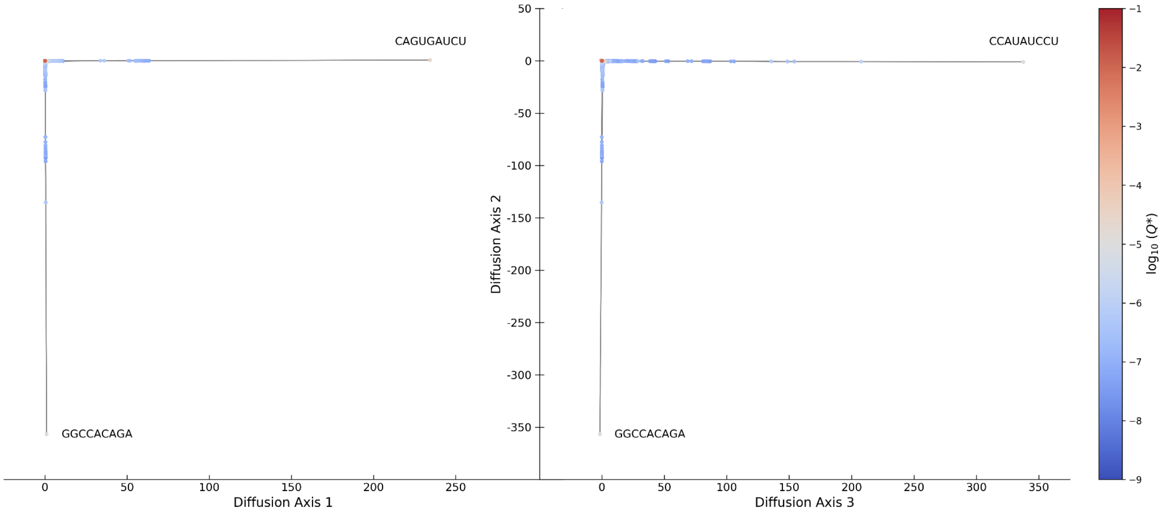


**Figure 3:** Visualization of the probability landscape $Q^*$ computed by SeqDEFT for human 5' splice sites with diffusion axis 1, 2, and 3. The RNA sequences at the extremes have unusual nucleotides at the +1 and +2 sites.

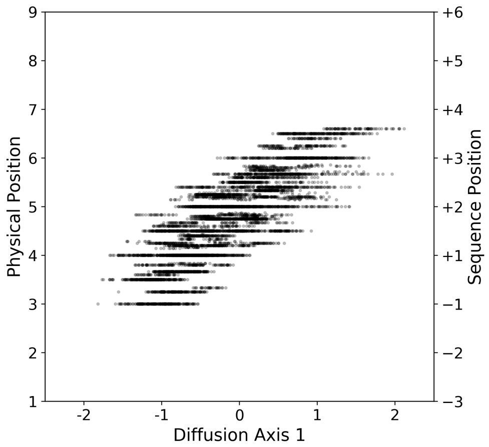


**Figure 4:** Average position of consensus nucleotides versus diffusion axis 1 for the RNA sequences in Figure 4 of the main text with values on diffusion axis 2 between from $-0.5$ to $0.5$ (i.e. the streak of consensus-like high probability sequences). Here we represent the sequence sites -3, -2, -1, +1, ..., +6 by ordinal numbers from 1 to 9. For each RNA sequence, its averaged physical position is computed by first finding the sites that match the canonical sequence CAGGUAAGU, then taking the average of the sites. For example, the sequence CAGGUUCAA matches the canonical sequence at sites -3, -2, -1, +1, and +2. The averaged physical position of this sequence is then equal to $(1+2+3+4+5)/5 = 3$. The right y-axis shows the corresponding positions in the original numbering system.

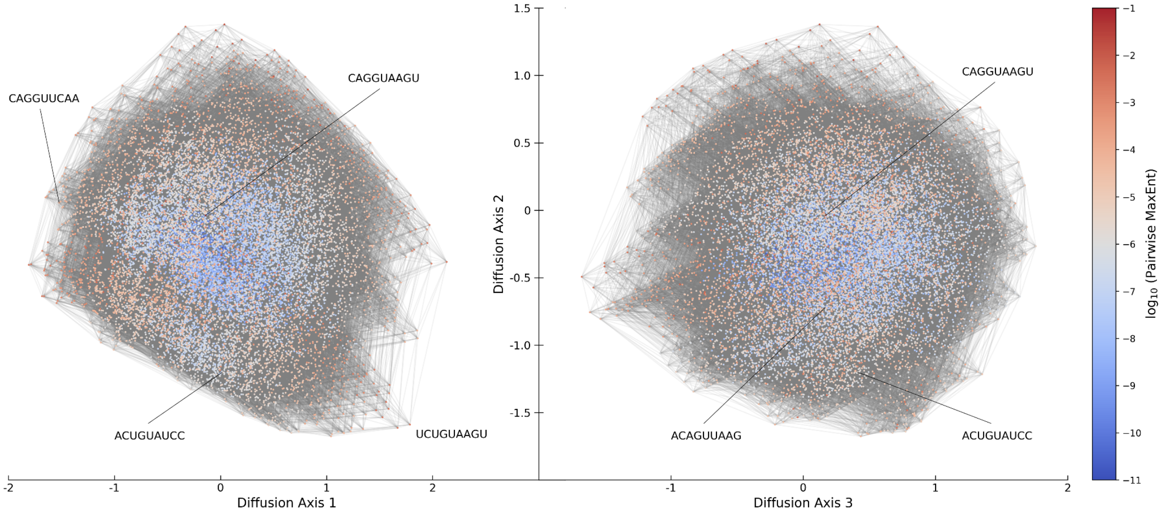


**Figure 5:** Visualization of the pairwise MaxEnt probability landscape for human 5' splice sites with diffusion axis 1, 2, and 3. The RNA sequences marked here are the same RNA sequences highlighted in the visualization of the optimal probability distribution $Q^*$ shown in Figure 4 of the main text.

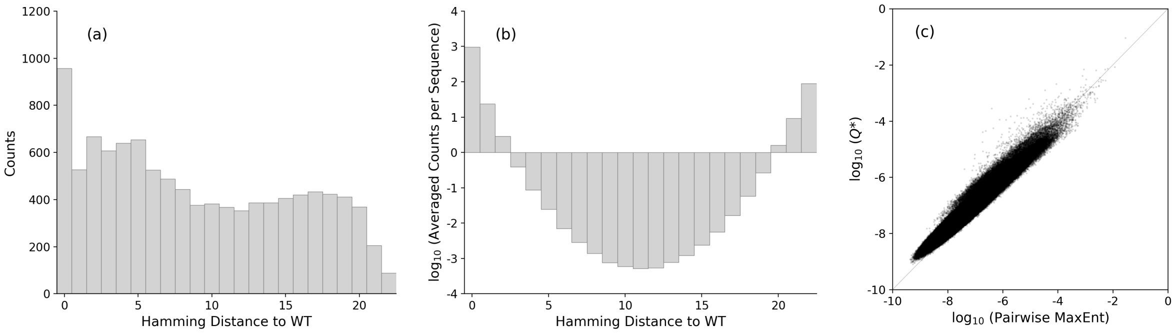


**Figure 6:** (a) Histogram of Hamming distance to the wild type (i.e., all the 22 chromosomes are normal) for karyotypes of human cancer. (b) Same as (a) with each count divided by the number of all possible sequences with certain Hamming distance to the wild type. (c) Comparison between the probability distribution $Q^*$ estimated by SeqDEFT and the probability distribution estimated by pairwise MaxEnt for the aneuploidy data.

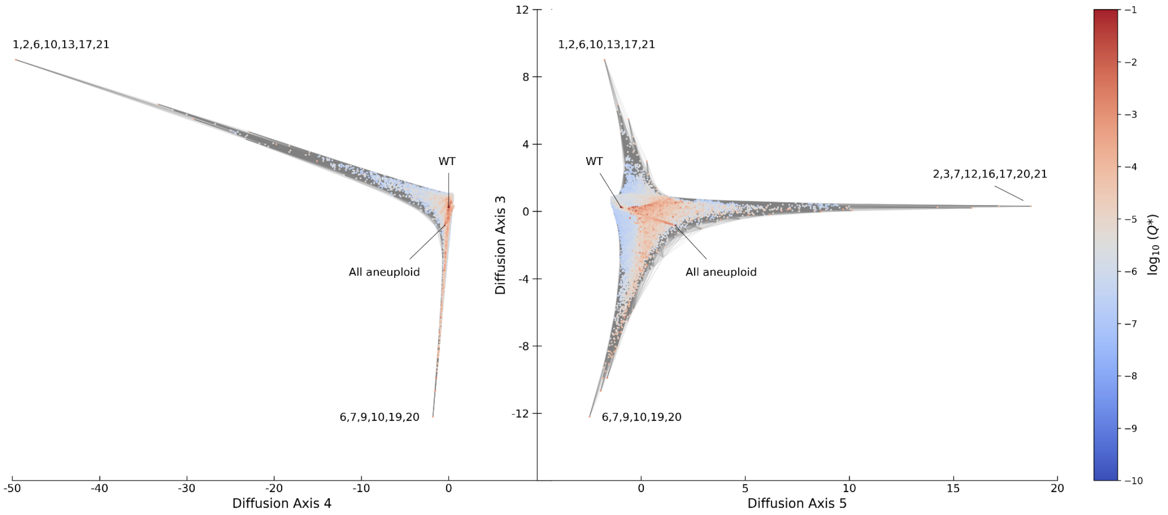


**Figure 7:** Visualization of the probability landscape $Q^*$ computed by SeqDEFT for karyotypes of human cancer with diffusion axis 3, 4, and 5. "WT" represents the wild type where all the 22 chromosomes are normal. The 3 subsets of chromosomes marked here are the same subsets that stand out in the visualization of $Q^*$ with diffusion axis 1, 2, and 3 shown in Figure 5 of the main text.

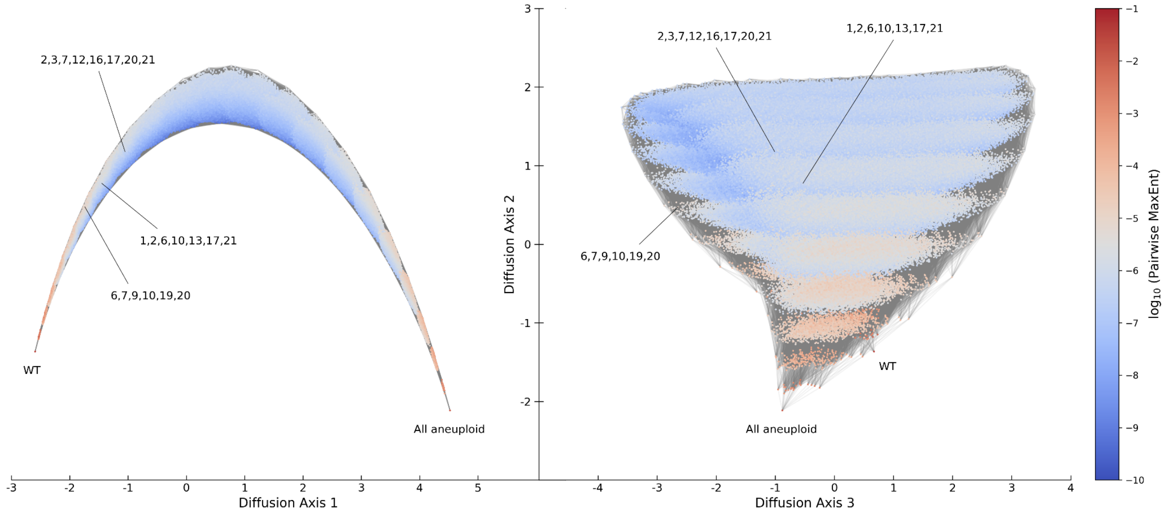


**Figure 8:** Visualization of the pairwise MaxEnt probability landscape for karyotypes of human cancer with diffusion axis 1, 2, and 3. "WT" represents the wild type where all the 22 chromosomes are normal. The 3 subsets of chromosomes marked here are the same subsets that stand out in the visualization of the optimal probability distribution $Q^*$ shown in Figure 5 of the main text.

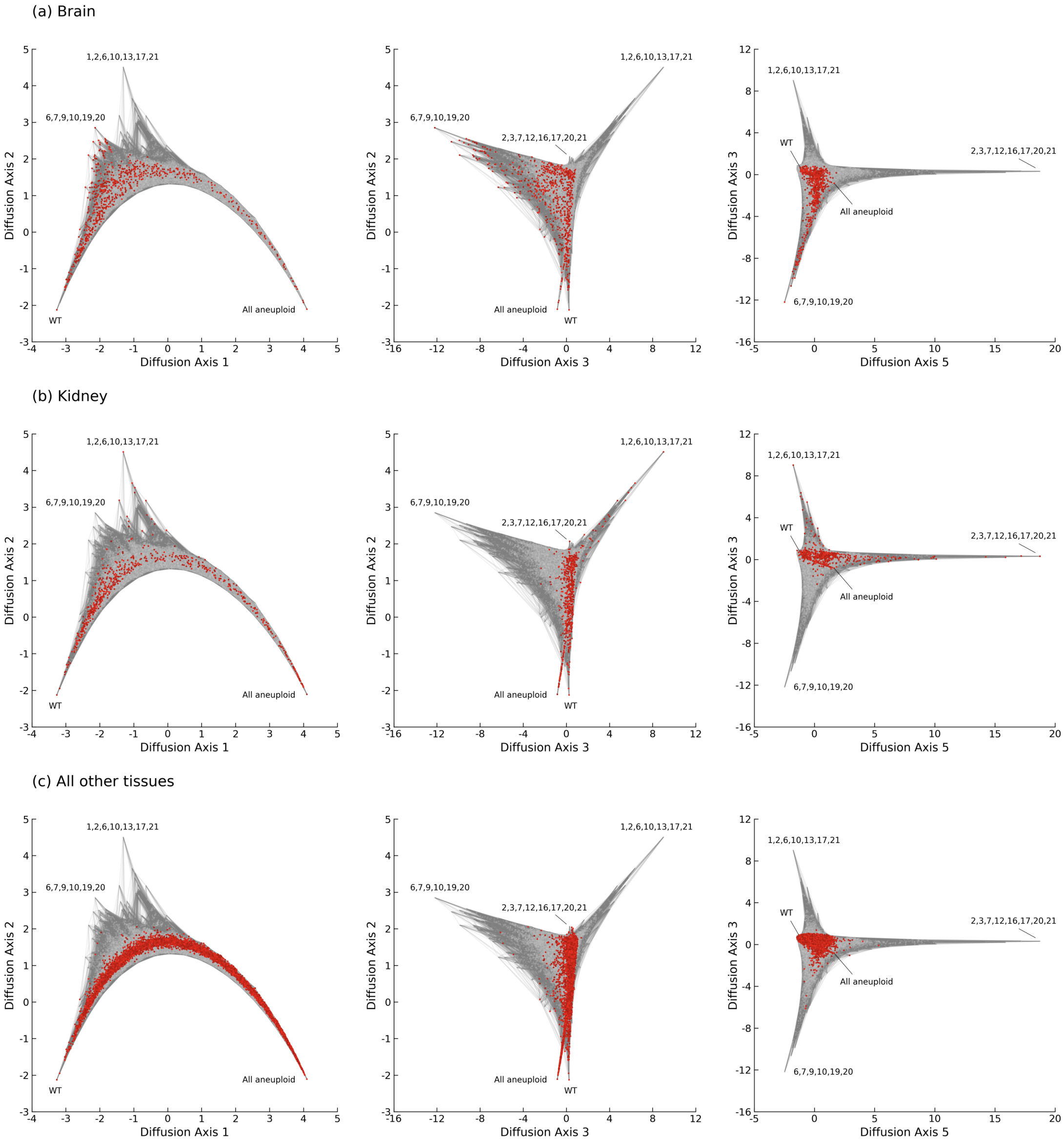


**Figure 9:** Visualizations of the probability landscape $Q^*$ computed by SeqDEFT for karyotypes of human cancer with 3 groups of tissues highlighted in red: (a) Brain, (b) Kidney, and (c) All other tissues. "WT" represents the wild type where all the 22 chromosomes are normal.

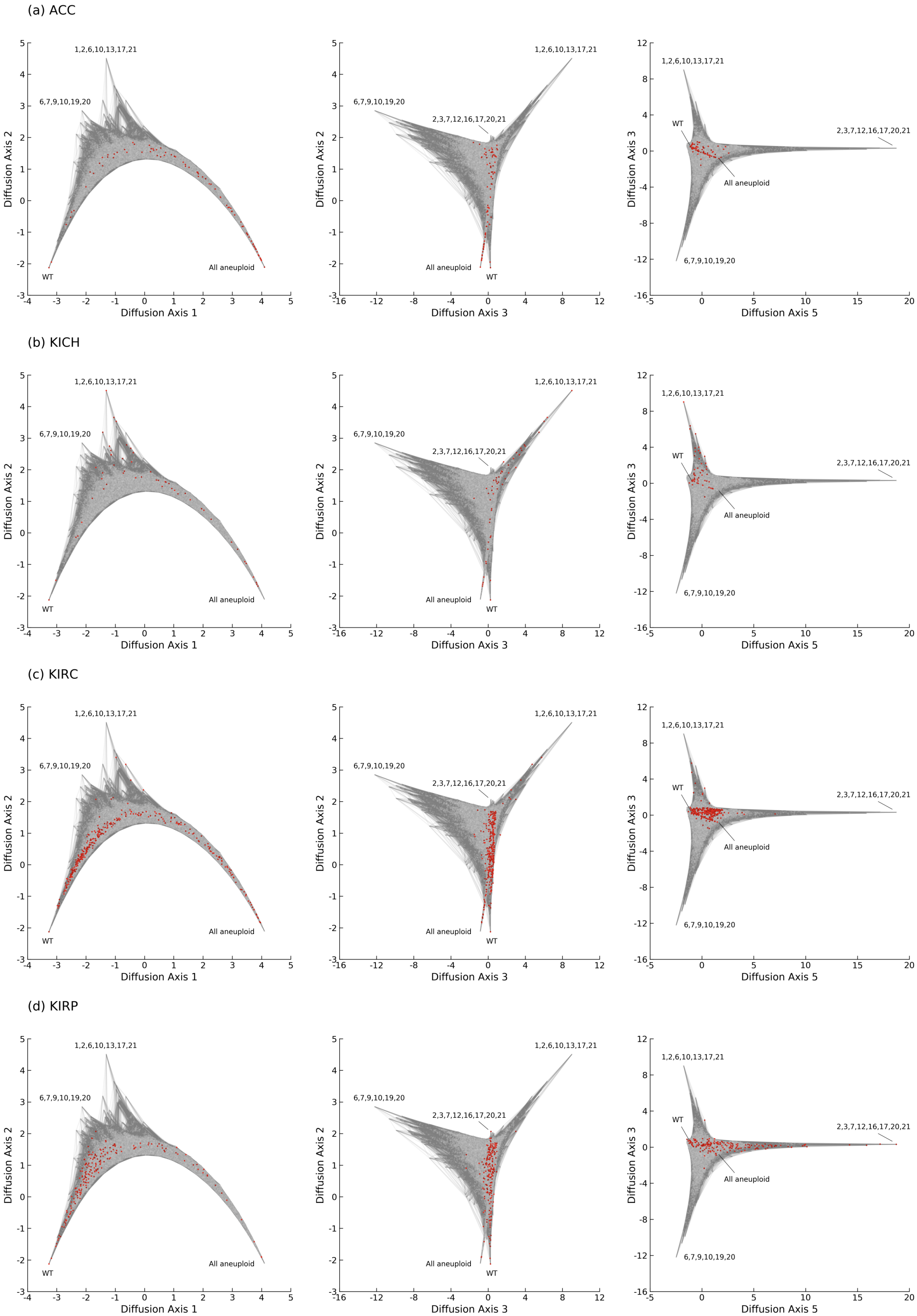


**Figure 10:** Same as Figure 9 but for the 4 subtypes of kidney-related cancer: (a) ACC (adrenocortical carcinoma), (b) KICH (kidney chromophobe), (c) KIRC (kidney renal clear cell carcinoma), and (d) KIRP (kidney renal papillary cell carcinoma).

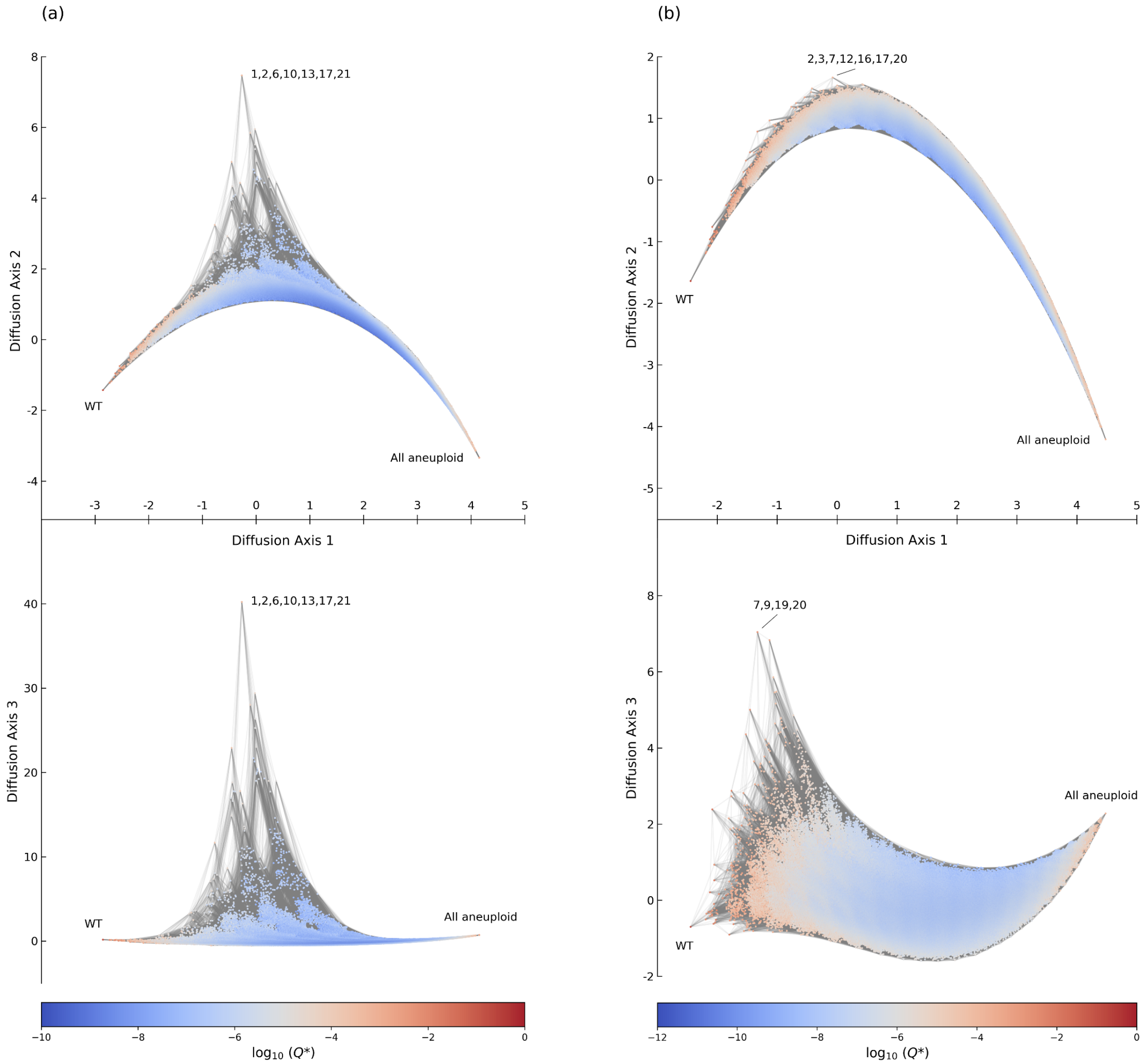


**Figure 11:** Visualization of the probability landscape $Q^*$ computed by SeqDEFT for karyotypes of human cancer with two different codings of the data: (a) 0 = normal or gain, 1 = deletion or complex, and (b) 0 = normal or deletion, 1 = gain or complex. "WT" represents the wild type where all the 22 chromosomes are in state 0, and "All aneuploid" means all the 22 chromosomes are in state 1. The peak in panel (a) corresponds to deletion in chromosomes 1, 2, 6, 10, 13, 17, 21, while the peaks in panel (b) correspond to gain in chromosomes 2, 3, 7, 12, 16, 17, 20 (top) and chromosomes 7, 9, 19, 20 (bottom). Thus, the peak in panels (a) and (b) are equivalent to those described in main text Figure 5, with the minor difference that the peak {6, 7, 9, 10, 19, 20} in Figure 5, which consists of a mixture of losses, gains, and complex aneuploidies in Figure 5, is represented here by only a subset of those chromosomes.

**Table 1:** Entropy $H$, effective number of sequences $2^H$, and percentage of effective sequences $2^H/G$ of the two datasets studied in this work computed with a number of probability distribution estimates: observed frequencies, SeqDEFT, pairwise MaxEnt, independent sites model, and uniform distribution. SeqDEFT estimates based on $a^*$, with a prior specified by $\Delta^{(2)}$ for human 5' splice sites and $\Delta^{(3)}$ for the binary encoding of cancer chromosomes as euploid vs. aneuploid.

| | Human 5' splice sites | | | Karyotypes of human cancer | | |
|---|---|---|---|---|---|---|
| | $H$ (bits) | $2^H$ | $2^H/G$ (%) | $H$ (bits) | $2^H$ | $2^H/G$ (%) |
| Observed frequencies | 9.91 | 960.84 | 0.37 | 11.68 | 3,284.82 | 0.08 |
| SeqDEFT posterior samples 95% CI | 9.99 ~ 10.02 | 1,019.55 ~ 1,035.12 | 0.39 ~ 0.39 | 15.85 ~ 16.13 | 59,018.56 ~ 71,655.46 | 1.41 ~ 1.71 |
| SeqDEFT MAP | 10.04 | 1,053.26 | 0.40 | 16.39 | 85,774.81 | 2.05 |
| Pairwise MaxEnt | 10.14 | 1,131.88 | 0.43 | 16.70 | 106,600.83 | 2.54 |
| Independent sites | 10.70 | 1,665.77 | 0.64 | 21.28 | 2,551,693.42 | 60.84 |
| Uniform distribution | 18.00 | 262,144.00 | 100.00 | 22.00 | 4,194,304.00 | 100.00 |

**Table 2:** Comparison between the field-theoretic methods for one-dimensional density estimation (DEFT) [1] and field-theoretic density estimation in sequence space (SeqDEFT).

| | DEFT | SeqDEFT |
|---|---|---|
| Space | One-dimensional continuous space | Discrete combinatorial space |
| Entity to estimate | Probability density | Probability |
| Practical dimension | Number of grid points | Number of all possible sequences |
| Smoothness measure | Derivatives (e.g., slope, curvature, . . . ) | Associations (e.g., odds, odds ratio, . . . ) |
| Solution at $a \to 0$ | Histogram | Observed frequency |
| Solution at $a \to \infty$ | Uniform, exponential, Gaussian, . . . | Constant, additive, pairwise, . . . |
| Quantities conserved along MAP curve | Moments (e.g., mean, variance, . . . ) | Marginal frequencies (e.g., 1-site, 2-site, . . . ) |
